# Massive invasion of organellar DNA drives nuclear genome evolution in *Toxoplasma*

**DOI:** 10.1101/2023.05.22.539837

**Authors:** Sivaranjani Namasivayam, Cheng Sun, Assiatu B Bah, Jenna Oberstaller, Edwin Pierre-Louis, Ronald Drew Etheridge, Cedric Feschotte, Ellen J. Pritham, Jessica C. Kissinger

## Abstract

*Toxoplasma gondii* is a zoonotic protist pathogen that infects up to 1/3 of the human population. This apicomplexan parasite contains three genome sequences: nuclear (63 Mb); plastid organellar, ptDNA (35 kb); and mitochondrial organellar, mtDNA (5.9 kb of non-repetitive sequence). We find that the nuclear genome contains a significant amount of NUMTs (nuclear DNA of mitochondrial origin) and NUPTs (nuclear DNA of plastid origin) that are continuously acquired and represent a significant source of intraspecific genetic variation. NUOT (nuclear DNA of organellar origin) accretion has generated 1.6% of the extant *T. gondii* ME49 nuclear genome; the highest fraction ever reported in any organism. NUOTs are primarily found in organisms that retain the non-homologous end-joining repair pathway. Significant movement of organellar DNA was experimentally captured via amplicon sequencing of a CRISPR-induced double-strand break in non-homologous end-joining repair competent, but not *ku80* mutant, *Toxoplasma* parasites. Comparisons with *Neospora caninum*, a species that diverged from *Toxoplasma* ∼28 MY ago, revealed that the movement and fixation of 5 NUMTs predates the split of the two genera. This unexpected level of NUMT conservation suggests evolutionary constraint for cellular function. Most NUMT insertions reside within (60%) or nearby genes (23% within 1.5 kb) and reporter assays indicate that some NUMTs have the ability to function as cis-regulatory elements modulating gene expression. Together these findings portray a role for organellar sequence insertion in dynamically shaping the genomic architecture and likely contributing to adaptation and phenotypic changes in this important human pathogen.

**Significance Statement:** This study reveals how DNA located in cellular compartments called organelles can be transferred to the nucleus of the cell and inserted into the nuclear genome of apicomplexan parasite *Toxoplasma*. Insertions alter the DNA sequence and may lead to significant changes in how genes function. Unexpectedly, we found that the human protist pathogen, *Toxoplasma gondii* and closely-related species have the largest observed organellar genome fragment content (>11,000 insertion comprising over 1 Mb of DNA) inserted into their nuclear genome sequence despite their compact 65 Mb nuclear genome. Insertions are occurring at a rate that makes them a significant mutational force that deserves further investigation when examining causes of adaptation and virulence of these parasites.

## Introduction

*Toxoplasma gondii* is a cosmopolitan apicomplexan protist parasite (Fig. 1A), capable of infecting nearly all warm-blooded animals including humans (1, 2). *T. gondii* causes toxoplasmosis, and the parasite infects as much as one third of the world’s human population with individual countries showing infection rates ranging from 10 to 80% (3). While the parasite rarely causes symptoms in healthy adults, it can lead to serious and even life-threatening illness in immunosuppressed or pregnant individuals (4, 5). Phylogenetic and population studies of *T. gondii* reveal an unusual population structure consisting of mostly clonal lineages that occasionally recombine (6–9) as well as the recent evolution of oral infectivity that permits the parasite to spread asexually via consumption of tissue cyst forms in addition to traditional transmission of sexually derived oocysts in the environment via a fecal/oral route (10). While it is clear that *Toxoplasma* and other coccidian parasites evolve and adapt rapidly, little is known about the source(s) of genomic variation driving adaptation in *Toxoplasma*. The ability of *Toxoplasma* to propagate asexually, combined with the observed low levels of sexual recombination in existing data and the lack of detectable mobile elements create real challenges for *Toxoplasma* with respect to sources of genomic variation. When contemplating genome evolution and the differences between strains and species it is common practice to examine single nucleotide polymorphisms (SNPs), insertion/deletion events (indels), rearrangements, horizontal and intracellular gene transfers and transposable element (TE) insertion or movement. The insertion of random fragments of organellar DNA is not high on the list of features examined as a major cause of phenotypic or genotypic diversity, but if the organism is a tissue coccidian like *Toxoplasma*, perhaps it should be.

**Figure 1.**
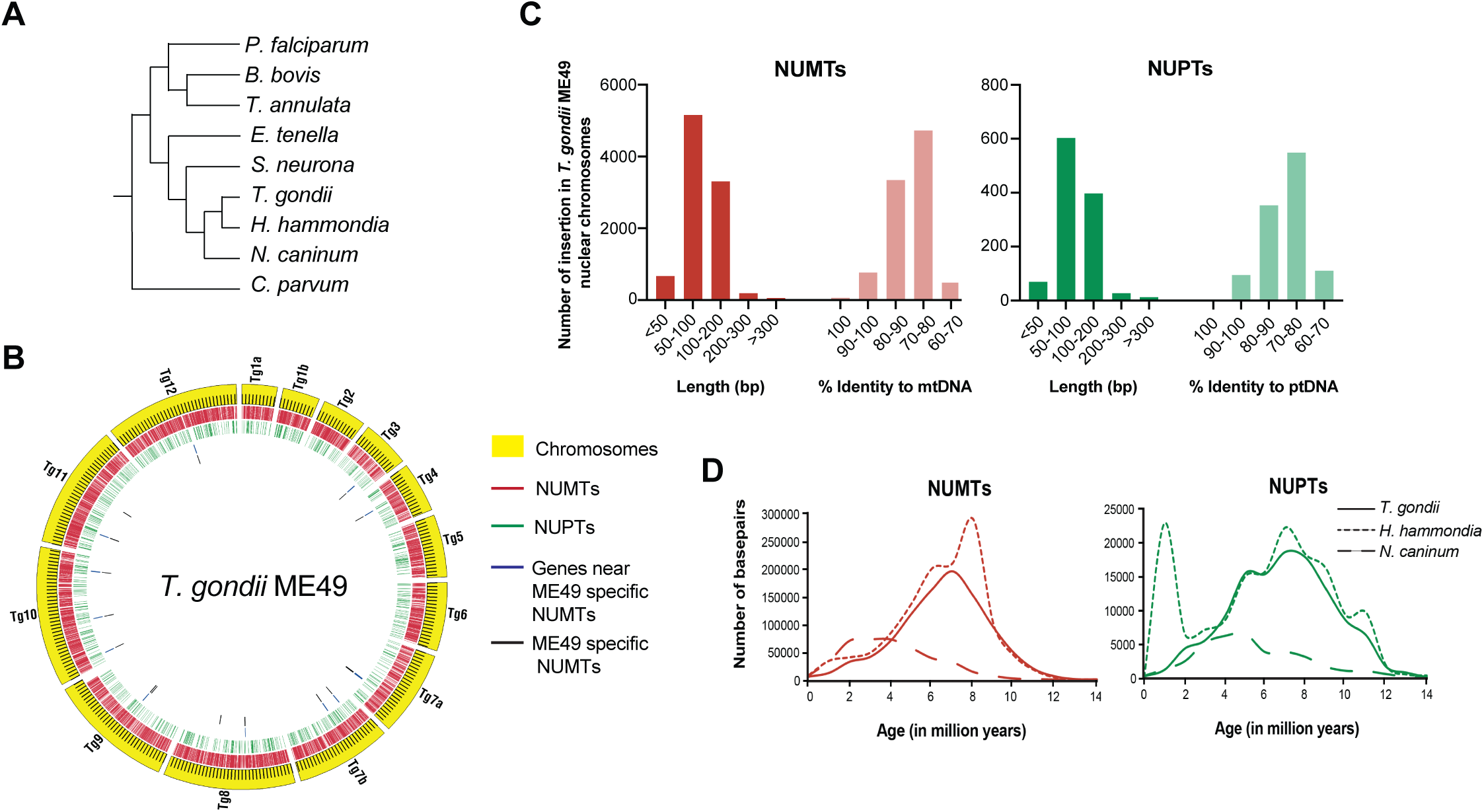
Characteristics of NUMTs and NUPTs in *T. gondii* ME49. *(A)* Cladogram of apicomplexan parasite relationships *(B)* Circos plot (Circos version 0.51) represents the distribution of NUMTs and NUPTs in the nuclear chromosomes of *T. gondii* ME49. The outer circle of yellow bands indicates the chromosomes as labeled and each tick on the band equals 100 kb. The ticks interior to the chromosomes are different features as indicated in the key. *(C)* The length and the percent identity of a NUMT and NUPT to mt and ptDNA respectively was calculated from RepeatMasker results and the distribution is plotted. *(D)* The age of each NUMT and NUPT was calculated based on the percent divergence from the mt and ptDNA of the corresponding species and the distribution of the age is plotted against the number of base pairs.

In eukaryotes, DNA of endosymbiotic organelles (mitochondria and chloroplasts) has been observed to be transferred to the nuclear genome (11). During the early phase of organelle evolution, this process resulted in a massive relocation of entire organellar genes to the nuclear genome (12). In many eukaryotes, the transfer of functional organellar genes appears rare or has ceased altogether (13, 14), and almost all recent transfers of mitochondrial (mtDNA) or plastid (ptDNA) DNA to the nuclear genome are gene or genome fragments that give rise to noncoding sequences, called NUMTs (nuclear integrants of mtDNA) and NUPTs (nuclear integrants of ptDNA). Organellar-to-nuclear DNA transfer has been reported for 85 fully sequenced eukaryotic genomes and fully characterized in 23 mammals (15). The results are indicative of the significant driving force NUOTs (nuclear DNA of organellar origin) provide for gene and genome innovation in eukaryotes (11) including humans (16) and other primates (17, 18). In *Saccharomyces cerevisiae*, 16 NUMTs were unexpectedly recovered in a double-strand break experiment (19) and the non-homologous end-joining repair pathway, NHEJ, was shown to be the mechanism responsible for NUMT integration (19, 20). Mechanistically, reactive oxygen species (ROS) can generate double-stranded mtDNA breaks (DSBs) and mtDNA fragments (21) and there are hypotheses and some evidence for how organellar DNA, in particular mtDNA may enter the host nucleus and be used to patch nuclear DSBs (22).

Nuclear genome sequences in the Apicomplexa are highly streamlined, ranging from 8 to 130 Mb, dynamic in structure and rapidly evolving (23, 24). Transposable elements (TEs), which generally constitute the most significant proportion of repetitive DNA in genomes and are known to be powerful agents for generating genomic plasticity (25, 26), are absent from most apicomplexans examined thus far (27) excepting the genera *Ascogregarina*, *Eimeria* and *Sarcocystis* (28, 29). Consequently, it is of interest to understand what factors contribute to the generation of genomic plasticity in apicomplexan parasites. While organellar-to-nuclear DNA transfer represents a significant driving force for genome innovation in eukaryotes, it has not been examined in apicomplexans (outside of nuclear-encoded, organellar-targeted proteins) (30, 31), and little is known about the evolutionary fate and consequence of these transferred DNA sequences.

Most apicomplexans contain two organelles, a mitochondrion and a plastid (pt) called the apicoplast that was acquired through the secondary endosymbiosis of a red alga (32–34). The ptDNA sequence (35 kb) is a circular/cruciform structure and is well conserved across apicomplexans (35). The mtDNA sequences generated thus far are variable and occur as linear monomers (∼6 kb) or concatemers with, or without, inverted repeats depending on the species (36, 37). Despite the genome topological differences, the apicomplexan mtDNA consistently encodes three cytochrome genes: cytochrome oxidase subunits I and III (*coxI, coxIII)* and cytochrome b (*cob*) and highly fragmented large and small subunit rRNA genes (LSU – SSUrRNA) (36, 37). Unlike the ptDNA sequence, the mtDNA sequence of *T. gondii* remained elusive until recently (36). It is structurally divergent and consists of 21 unique sequence blocks (SBs) that represent highly fragmented protein coding and rRNA genes (36). These sequence blocks are highly redundant, as was observed in dinoflagellates (38, 39). The presence of NUMTs in *Toxoplasma* was first reported as REP elements in 1991 (40), but they have not been studied since. The availability of genome sequences for numerous *Toxoplasma* strains as well as organellar DNA sequences, permitted a systematic examination of NUOTs in *T*. *gondii* and its relatives to shed light on their evolutionary dynamics and impact on genome innovation.

## Results

### Exceptional density of NUMTs and NUPTs in *T. gondii*

To identify insertions of organellar DNA fragments in the nuclear genome of *T. gondii* ME49 we applied the homology-based program Repeat Masker with organellar genomes as libraries. Only the 14 nuclear chromosomal assemblies and not all available contigs were queried for organellar insertions to obtain reliable and conservative estimates. The 21 conserved mtDNA sequence blocks (36) as well as full-length sequences of the mitochondrial protein-coding genes, *coxI*, *coxIII* and *cob* were included in the repeat library to minimize overestimation of the number of NUMTs. The 35 kb apicoplast genome sequence of *T. gondii* was utilized as the repeat library to identify NUPTs. These analyses revealed that 1.43% (895,312 bp) and 0.18% (112,433 bp) of the nuclear genome of *T. gondii* MR49 is composed of NUMTs and NUPTs respectively (Table 1, Fig. 1B, Dataset S1). To our knowledge, this percentage of nuclear genome occupied by NUOTs represents the largest ever reported for any eukaryote (Table 1). Indeed, the NUMT insertions in *T. gondii* are almost 6 times higher than the current overall record holder, the pathogenic fungus *Ustilago maydis* (0.286%) (41) (Table 1).

**Table 1.**
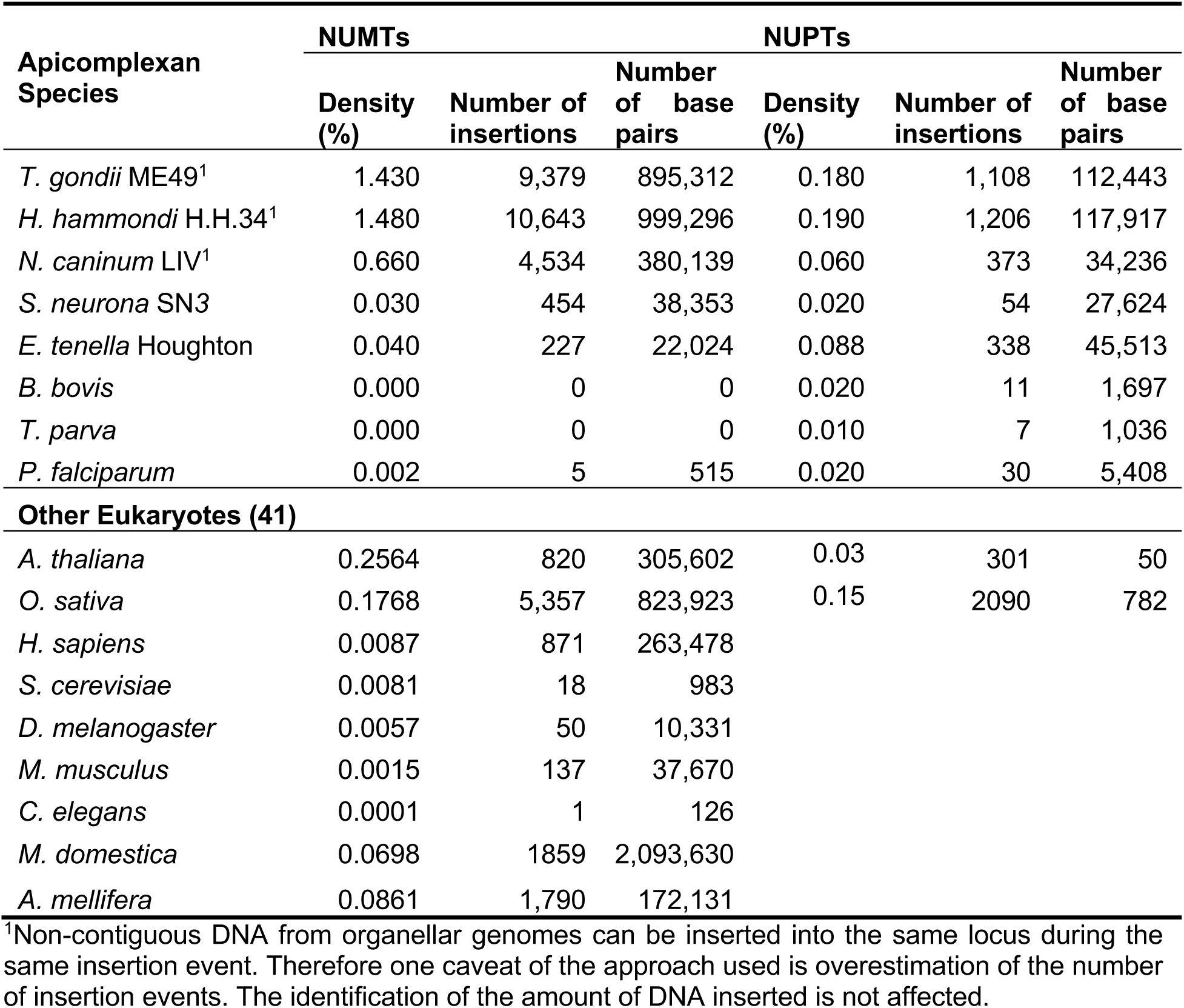
NUMTs and NUPTs detected in apicomplexan other eukaryotic genome sequences

Most (>90%) of the NUOTs identified in *T. gondii* are 50-200 bp in length and display <90% sequence identity to their respective organellar DNA counterparts indicating most insertions have accumulated mutations after nuclear acquisition (Fig. 1C). The largest NUMT is a single 3,369 bp sequence on Chr 10 that does not contain any complete coding sequences but retains 99% nucleotide identity to mtDNA sequence blocks suggesting it is a recent insertion (Table S1). NUMTs derive from all 21 mtDNA SBs including coding and non-coding regions (Table S2). Sequence blocks M, E, V and K are the most enriched each comprising at least 0.1 % of the nuclear genome (Table S2). Similarly, NUPTs arise from nearly all regions of the apicoplast genome, with the region between coordinates 9600 to 12000 being the least represented region in NUPTs (Dataset S1).

We next examined the location and distribution of NUOTs in the *T. gondii* nuclear genome. Approximately 60% of the identified NUOTs map to genic regions (introns and exons) with the vast majority located in introns and the rest almost exclusively in UTRs (Table S3). The other 40% of NUOTs are located in intergenic regions, and ∼23% occur within the 1 kb flanking region of genes. Thus, ∼85% of *T. gondii* NUOTs reside within or near genes. This distribution is not unexpected given the highly compact nature of the *T. gondii* genome where genes occupy ∼70% of the nuclear sequence.

To determine if NUOTs arose due to independent insertion events or were segmentally duplicated post-insertion, we examined the flanking regions of NUOTs as duplication would likely involve flanking nuclear sequence. We considered NUOTs to be the product of a nuclear segmental duplication event if the NUOT along with 100 bp of its flanking regions was detected more than once. Using this criterion, 105 NUMTs and 13 NUPTs were identified as segmentally duplicated two to ten times in the ME49 nuclear genome sequence (Table S4). Since <1% of the NUOTs were duplicated post-insertion suggesting, most NUOTs were independent insertion events. Together, these findings demonstrate that the large proportion of NUOTs observed in the nuclear genome of *T. gondii* arose primarily through independent insertion events that are generally <200 bp in length and display varying levels of sequence degeneration relative to their organellar DNA counterparts.

### Strain-specific NUMTs indicate recent insertions and frequent turnover

To determine the age distribution of the NUOTs, age-related decay was determined using T = K/2μ where μ is the intron mutation rate for *T. gondii* (10) and K, the genetic distance, was determined with the Jukes-Cantor model based on the difference between the NUOT and organellar sequences (see Methods). This analysis revealed a continuous influx of NUOT assimilations with a NUMT accumulation peak approximately 7 million years ago (Fig. 1D). While NUOT assimilation appears to be a continual process, most insertions occurred prior to the last 1 million years. The presence of the NUOTs that are 100% identical to organellar DNA indicate that the process is ongoing.

To identify recent insertions or deletions we compared NUMTs in the nuclear genome of different *T. gondii* strains. We restricted this investigation to NUMTs, since the number of NUMTs is higher than NUPTs. We compared the NUMTs identified in *T. gondii* ME49 to those identified in strains, GT1 and VEG, both of which have chromosomal level nuclear genome assemblies. We identified a total of 32 strain-specific NUMTs: 12 in ME49, 13 in GT1 and 7 in VEG (Table S5). A few of these strain-specific NUMTs were experimentally verified via PCR (Fig. S1, Table S6). 26 of the 32 strain-specific NUMTs were found in or near annotated genes, suggesting that their presence may have the potential to affect host gene activity (Table S5).

To ascertain if the strain-specific NUMTs arose via either a novel insertion or deletion event, we determined sequence boundaries and performed a three-way genome comparison of the regions in question. A NUMT was considered a novel insertion if it was precisely missing from two of the three strains and displayed high sequence identity (>95%) when compared to the mtDNA sequence (Fig. 2A). If a NUMT was missing in only one of the three strains and the NUMT in the 2 strains displayed high sequence divergence (>10%) when compared to the mtDNA, then we concluded that this NUMT was most likely deleted in the third strain (Fig. 2B). Using such criteria, 3 NUMTs were confidently identified as novel insertions, and 13 were inferred to be deletions and many of these are partial deletions (Table S5). These results suggest that NUMTs are dynamic and can lead to strain differences in *Toxoplasma*.

**Figure 2.**
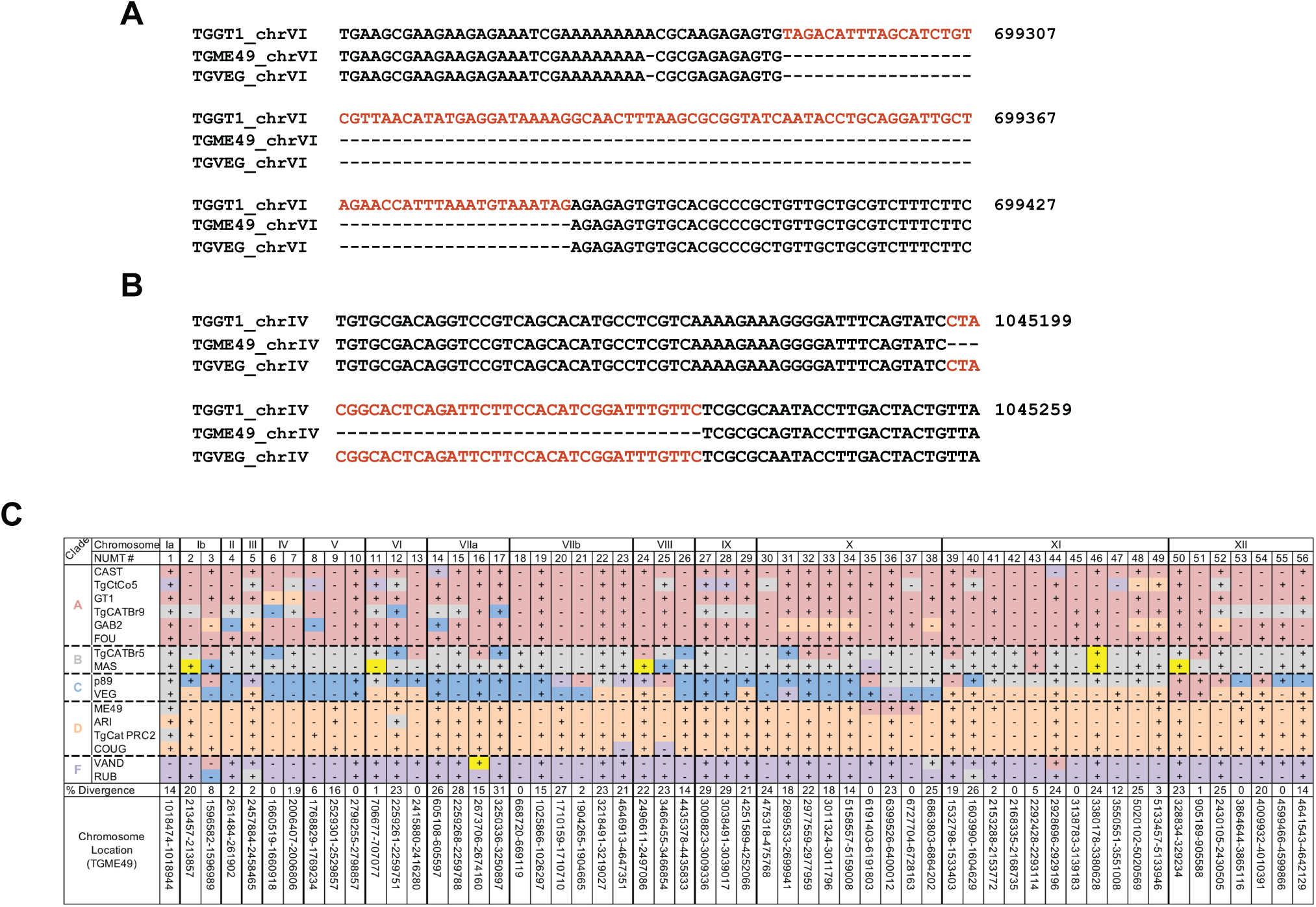
Strain-specific presence and absence of NUMTs in *T. gondii*. *(A, B)* A likely strain-specific insertion in *T. gondii* (TG) GT1 *(A)* and deletion in ME49 *(B)* are shown. A multiple sequence alignment of these loci was performed for strains GT1, VEG and ME49 and the co-ordinates of the loci are indicated with reference with GT1 chromosome VI. The NUMT and flanking nuclear sequence are indicated in red and black respectively. *(C)* NUMTs in 16 *T. gondii* strains were compared to identify a strain-specific presence and absence. Strains are grouped and colored by clades as previously identified (7). The 14 chromosomes divided by black lines are represented on the x-axis. NUMTs that show differential presence/absence in the 16 *T. gondii* strains are numbered and ordered based on their location in ME49. The coordinates of each NUMT plus 200 bp flanking region on the ME49 chromosomes are shown at the bottom. The 200 bp regions were included to provide a location for NUMTs missing in ME49. The percent divergence of the NUMT from the mtDNA sequence is indicated above the coordinates. In the matrix, “+” and “-” indicate presence and absence of a NUMT in a particular strain. Each cell is colored based on local clade ancestry of that location (7) although, it does not imply that location originated from that clade. Yellow cells indicate clade E ancestry and were not included the NUMT analysis as an assembled genome sequence for a strain from this clade was not available

To further explore the differential presence or absence of NUMTs in *T. gondii* we examined 13 additional strains with assembled, but less complete genome sequences. All of the strains displayed a similar NUMT content (Table S7). Overall, we identified 8,271 ancestral NUMTs that are present in all 16 strains and 56 NUMTs that show differential presence or absence among these strains (Fig. 2C, Table S7). These differentially present NUMTs are in varying states of decay, ranging from 0%-31% divergence from the mtDNA sequence. No strain shows a preferential loss or gain/retention of NUMTs. MAS has the largest number of differential NUMTs at 36 out of 56 and VEG the fewest (25/56) (Table S8).

The sites of insertion or deletion for the 56 differentially present NUMTs are mostly conserved among the strains. Thus, it is likely that these NUMTs were inserted in an ancestor and strains which evolved from this ancestor inherited the NUMT. For example, NUMTs 18 and 42 (100% identical to the mtDNA) are present only in strains MAS and TgCATBr5 and these strains are clustered in Clade B (Fig. 2C) (7). NUMTs also appear to be inherited via recombination with a strain that does not contain a given NUMT. For example, NUMTs 13, 35 and 37 that display 100% identity to the mtDNA are each present in only two strains (p89 and VAND, TgCtCo5 and VAND and GAB2-2007 and ME49 respectively) suggesting inheritance of the NUMT via recombination, or, less likely, independent insertion of the same NUMT at the same location in different strains. These pairs of strains are not members of the same clade and other clade members do not contain these NUMTs (Fig. 2C). Genome analysis of *T. gondii* has demonstrated that the parasites undergo sexual recombination at a higher frequency than previously thought (7). Our finding that NUMT presence or absence does not follow clades is consistent with this observation.

Finally, to examine potential functional consequences of the 56 differentially present NUMTs across the 16 *T. gondii* strains, we identified their insertion location with respect to genes. 8 are located in exons and 31 are located in introns. 11 NUMTs are located in flanking regions or UTR, the remaining 6 are in intergenic regions outside of the 1 kb upstream and downstream flanking regions (Table S8). None are located in a protein coding region. We conducted a gene-ontology (GO) enrichment analysis with the 50 genes whose exons, introns or flanking regions contained NUMTs. Genes involved in microtubule-based processes and movement (10 genes) and nucleotide metabolic or biosynthetic processes (10 genes) are enriched 5-20 fold (Table S9). These pathways and processes are core survival and virulence pathways as *T. gondii* acquires purines from its host via salvage rather than *de novo* synthesis (42). It is possible that NUMTs in these regions may affect transcription at these loci. However, transcriptome data are only available for 3/16 strains analyzed here and only for tachyzoites.

### NUOTs are drivers of genome evolution in coccidian parasites

To determine if the high NUOT density is a phenomenon specific to *T. gondii*, we investigated the nuclear genome sequences of its closest coccidian relatives *Hammondia hammondi* and *Neospora caninum* (Fig. 1A). *H. hammondi* contains NUMT and NUPT densities of 1.48% and 0.19% respectively, a level higher than that observed in *T. gondii*. However, because the *H. hammondi* genome sequence assembly is highly fragmented the NUOT densities are only estimates. The NUOT content in *N. caninum’s* nuclear genome is substantially lower than *T. gondii* at 0.66% for NUMTs and 0.06% for NUPTs, however, these densities are still higher than those observed in other eukaryotes (Table 1). By contrast, the distantly related coccidian species, *Sarcocystis neurona* and *Eimeria tenella* display greatly reduced NUMT densities at 0.03 and 0.04% respectively, relative to *T. gondii* and *N. caninum* but their NUPT densities, especially in the case of *E. tenella*, are only slightly reduced relative to *T. gondii* (Table 1). Examination of the apicomplexan nuclear genome sequences of species outside of the Coccidia revealed a paucity of NUOTs. *Plasmodium falciparum* 3D7 contains only five NUMTs whereas we were unable to detect any such insertions in *Babesia bovis* and *Theliera parva* (Table 1). Thus, an unusually high NUOT density is a feature of coccidian parasites and is more pronounced in species closely related to *T. gondii*.

Next, we examined the characteristics of the NUOTs in *N. caninum* and *H. hammondi*. Similar to *T. gondii*, we found that most NUMTs are 50-200 bp in length in both species (Figs. S2, S3). Greater than 90% of the NUOTs display less than 80% identity to the mt/ptDNA in *H. hammondi* as is seen in *T. gondii*. Interestingly, the NUOTs in *N. caninum* appear younger (Fig. 1D Figs. S2, S3) suggesting either more recent insertions, a different mutation rate or a more rapid turnover rate in this species. NUPT’s in *H. hammondi*, unlike the other coccidian species show two distinct age populations, one quite recent (Fig. S3) but this may be an artifact related to contaminating apicoplast sequences in the genome assembly. These results reveal that the species most closely related to *T. gondii*, also have a very high NUOT density but that as you move to more distant coccidian species, NUOTs decrease. In apicomplexans outside of the coccidia, NOUTs are mostly non-existent.

Since NUMTs display an overall rapid turnover and decay based on comparisons between *T. gondii* and *N. caninum* and within *T. gondii* strains, we hypothesized that evolutionary conserved NUMTs might be under functional constraint, indicating cooption for cellular function. To identify such coopted candidates, we compared the NUMTs between *T. gondii* and *N. caninum*, two species that diverged ∼28 MYA (43), and identified five orthologous NUMTs (Table S10). The orthologous NUMT residing in the upstream region of the U6 snRNA-associated Sm family protein encoding gene (TGME49_286560) was the most conserved NUMT, thus we selected this NUMT, named NUMT[Sm]. NUMT[Sm] is 28.8% divergent from mtDNA SB U which encodes portions of LSU and SSU rRNA fragments. Next we selected a younger *T. gondii* specific NUMT, called NUMT[Myo] located in the upstream region of a putative myosin heavy chain gene (TGME49_254850) of all examined *T. gondii* strains. NUMT[Myo] is 16.7% divergent from mtDNA SB F which encodes RNA 10 (Fig. S3)(36). The upstream regions of both genes show evidence of active transcription in *T. gondii* based on H3K9ac acetylation and H3K4 tri-methylation marks, ToxoDB.org (44)(Fig. S4). The upstream regions of both *T. gondii* genes were cloned and tested for their ability to drive luciferase gene expression in transient *in vitro* reporter assays in *T. gondii* cultured parasites. For each promoter we generated luciferase reporter constructs where the upstream region was wild-type or with the identified NUMT removed (Fig. 3A, B). Deletion of the 70 bp NUMT[Sm] significantly decreased promoter activity whereas deletion of the 66 bp NUMT[Myo] dramatically increased promoter activity (Fig. 3C, D). These results suggest that NUMTs can carry, or can evolve into cis-regulatory elements capable of modulating gene expression. Taken together, these data demonstrate that NUMTs undergo rapid turnover and that evolutionarily conserved insertions may have been retained due to functional constraints for regulatory function.

**Figure 3.**
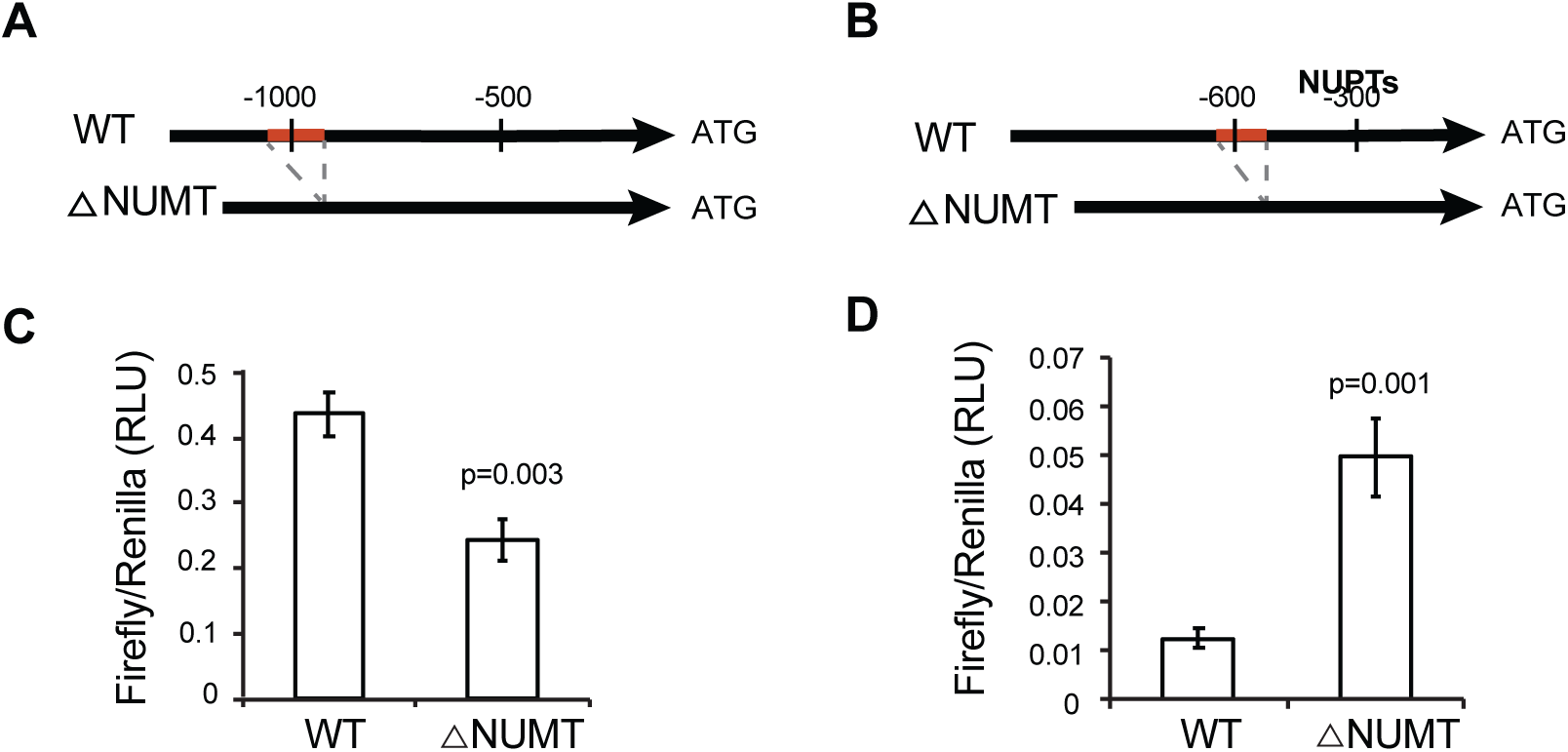
Cis-regulatory activity of two selected NUMTs. *(A, B)* The structure of WT and NUMT deletion promoter constructs for U6 snRNA-associated Sm family protein gene TGME49_286560 *(A)* and myosin heavy chain gene TGME49_254850 *(B)*. Nucleotide positions are referenced with respect to the start of translation (+1) of host gene and the red region indicates the position of the NUMT (Fig. S3). *(C, D)* Reporter assay results for the promoter of Sm-like protein *(C)* and mysoin heavy chain *(D)* genes. The graphs depict luciferase activity as ratios of Firefly:Renilla activity in relative luciferase units (RLU) from the different constructs containing either WT or mutagenized promoter. All luciferase readings are relative to an internal control (α-tubulin-Renilla). Error bars represent standard error calculated across the means of six independent electroporations. P-values were calculated using Student’s *t*-test.

### NUOT acquisition involves non-homologous end joining repair

It has been demonstrated that organellar DNA fragments can become integrated into the nuclear genome during the process of double-strand break (DSB) repair via a mechanism called non-homologous end joining (NHEJ) (19, 20). NHEJ can repair DSBs without a template or sequence homology but it often results in small deletions and in some cases insertion of short DNA sequences (45). This mechanism of integration of DNA fragments, usually NUOTs but also transposable elements was first demonstrated in *Saccharomyces cerevisiae* and subsequently proven in mammalian cells and plants (19, 20, 46, 47). In *T. gondii*, it has been observed that DNA is readily integrated into the nuclear genome at random sites via NHEJ (48). Indeed, it was necessary to knock-out the gene encoding a critical NHEJ protein, Ku80, in order to increase the efficiency of homologous recombination in this species (48). Interestingly, apicomplexans that lack the NHEJ machinery also contain very few NUOTs (Fig. 4A, Table 1), suggesting that NHEJ is a prominent mechanism underlying NUOT integration in coccidian species.

**Figure 4.**
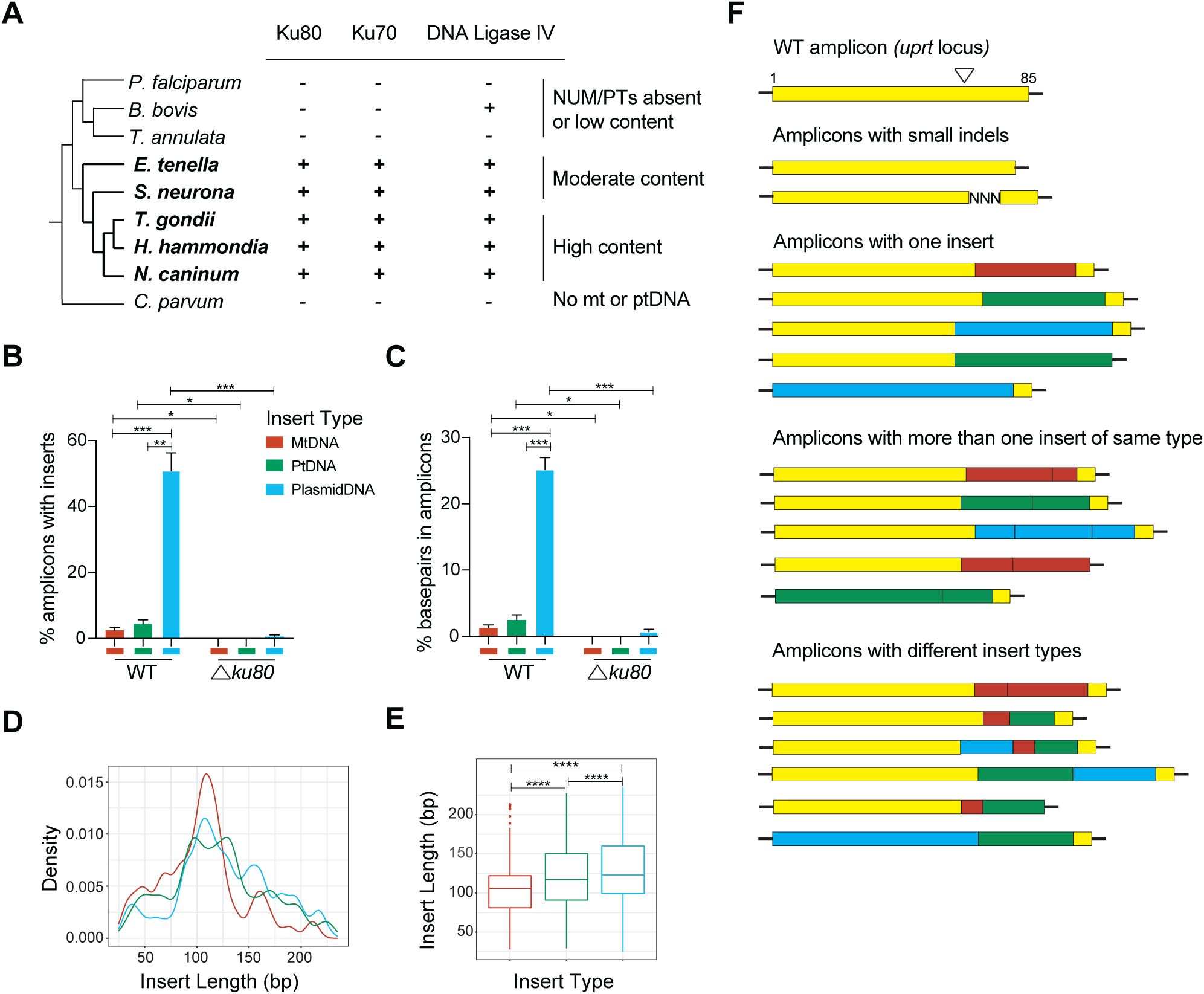
Capture of DNA insertions at CRISPR/Cas9-directed double-strand break at *uprt locu*s in wildtype and Δ*ku80* parasites. Utilizing the CRISPR-Cas9 system, double-strand breaks were induced at the *uprt* locus in wildtype *T. gondii* and Δ*ku80* parasites. The locus was then amplified and deep-sequenced to evaluate the extrachromosomal DNA insertions (mtDNA, ptDNA, CRISPR-Cas9 plasmid DNA) at the DSB site. *(A)* Apicomlexan cladogram with presence (+) or absence (-) of key NHEJ pathway genes and NUOT content in each species indicated. The coccidian branch is shown in bold. *(B, C)* The percent of amplicons with insertions from each DNA type *(B)* and the percentage of base pairs from each type present in the amplicons *(C)* are plotted for WT and Δ*ku80* parasites. The amplicons analyzed comprise all read pairs that could be merged as well as unmerged reads. Error bars represent standard error of means of three replicates. Statistical significance was calculated using Student’s *t*-test (* *p < 0.05*, ** *p < 0.01*, *** *p < 0.001*). *(D, E)* Distribution of the insert length for each DNA type is plotted as a proportion *(D)* and as box plots *(E)*. The data were generated from merged read pairs for one WT replicate. Color key is as shown in *(B).* Statistical significance was calculated using Student’s *t*-test (**** *p < 0.0001*). *(F)* Schematic representation of the modifications observed in the amplicon reads for WT parasites. The numbers under ‘WT amplicon’ indicate the length of the amplicon with the inverted triangle denoting the approximate site of the double strand break. The type of DNA inserted is indicated in the key in *(A).* Not all combinations of insertion patterns observed in the reads are represented. Examples of reads with the kind of modifications shown in this schematic are provided in Table S12. Amplicons with WT sequence (yellow) on both ends represent reads where the pairs could be merged. Numerous unmerged read with similar insertion patterns as shown in this figure exist but are not drawn here.

To experimentally test if NHEJ drives NUOT integration we used the CRISPR-Cas9 system to generate targeted DSBs in the genome of a *T. gondii* RH wildtype strain as well as a *ku80* knock-out mutant strain, Δ*ku80* (49, 50). We chose the *uprt* locus as the target for the DSB since disruption of this gene leads to 5-fluorodeoxyribose (FUDR) resistance thus facilitating efficient selection for improper repair of a CRISPR-Cas9 guide-directed cut (49). Resistant parasites were recovered, DNA was extracted, the *uprt* locus was amplified and the amplicons were deep-sequenced using Illumina paired-end sequencing (Table S11). Processed reads were then analyzed to compare the number of insertions between wildtype and Δ*ku80* parasites and specifically for insertions of extra-chromosomal DNA, including NUOTs and CRISPR-Cas9 plasmid DNA. It should be noted that nuclear DNA was also observed to be inserted at the DSB site but was not further investigated.

Overall, we observed that the number of unique reads obtained from WT parasites was seven times higher than that obtained from the Δ*ku80* strain (Table S11). Less than 1% of the reads from the Δ*ku80* parasites contained inserts of extrachromosomal DNA, whereas such insertions were observed in ∼58% of the amplicons from wildtype *T. gondii* (Fig. 4B). Analysis of the source of DNA inserted at the DSB site revealed the transfected CRISPR-Cas9 plasmid was by far the most common source of inserted extra-chromosomal DNA both in terms of number of reads and number of base pairs inserted (Fig. 4B, C). We expected the majority of the inserts to be from plasmid DNA since it is readily available for repair following transfection. However, both mtDNA and ptDNA sequences were recovered as well, but surprisingly ptDNA insertions were slightly more common than mtDNA. This finding is in contrast to the observed higher levels of NUMTs versus NUPTs in the nuclear chromosomes of tissue coccidia (Table 1). Importantly, DNA capture was drastically reduced in the Δ*ku80* strain with no organellar DNA recovered at the DSB repair site (Fig. 4B, C).

We next compared insert length. Since only 150 bp from each end of the amplicon was sequenced, this analysis is limited to detecting a maximum insert length of ∼250 bp. We observed that a large proportion of inserts from mtDNA were shorter (∼110 bp) than those from ptDNA and plasmid DNA with the latter being the longest (Fig. 4 D, E). Inserts as long as 250 bp are detected from all three DNA sources. As is typically seen in NHEJ mediated repair, we detected insertion or deletion of a few base pairs at the repair site. We also observed simple insertions of one continuous stretch of DNA and a number of complex insertions where non-contiguous stretches of DNA from the same or multiple DNA sources were observed (Fig. 4F, Table S12). The insertions derived from all parts of the mt, pt or plasmid DNA and are of varying lengths (Table S12). These data indicate that DSB repair via NHEJ is the likely mechanism underlying NUOT integration in *T. gondii.* The properties of the observed insertions in an artificially induced DSB are similar to those observed in naturally acquired NUOTs with the exception that NUPTs were observed more often than NUMTs.

## Discussion

DNA of organellar origin,NUOTs, can also be an important source of genetic variation. For example, a recent study of NUMTs in 66,038 human genome sequences revealed a complex and dynamic landscape with *de novo* NUMTs appearing in the germline once in every 10^4^ births and somatic insertions once in every 10^3^ cancer cells highlighting the ongoing formation and mutagenic potential of NUMTs in our species (16).

Our study shows that in the *Toxoplasmatinae*, which lack mobile elements, NUOTs are remarkably common and have been an important force in generating variation and evolution. To our knowledge, the *T. gondii* nuclear genome harbors the highest density of NUOTs ever reported at 1.61%. This percentage is more than 5-fold higher than the previous record holders *Arabidopsis thaliana* (0.25%) and *Ustilago maydis* (0.28%) (41).

The majority of the NUOTs in *T. gondii* arose from independent insertions rather than segmental duplications of previous insertions (51). However, unlike humans where NUMT insertions are concentrated in pericentromeric and subtelomeric regions (52). NUOTs in *T. gondii* and *N. caninum* occur evenly throughout the chromosome. A previous study reporting the presence of NUMTs in *T. gondii* found inverted repeats flanking the insertions (40). We have determined that the inverted repeat sequences are derived from mtDNA sequence block J, which is part of the *coxIII* gene. In our systematic catalog of 9379 NUMTs in the *T. gondii* genome, we rarely see inverted repeats flanking the NUMTs and SB J does not make up a higher proportion of the observed NUMT sequence, suggesting it is unlikely to be involved in propagating the NUMTs as had been suggested. The mtDNA of *T. gondii* is full of redundant sequence blocks often in alternative orientations within the same molecule (36). It should be noted that the high level of NUMTs in the nuclear genome, especially those with high similarity to the mtDNA was a significant confounding factor for the elucidation of the bizarre mtDNAs of *T. gondii*, *H. hammondi* and *N. caninum* as they made it impossible to use nucleic acid hybridization, PCR and genome assembly software (36).

Molecular and bioinformatic studies performed in yeast, tobacco and human have shown that integration of organellar DNA fragments occur during illegitimate repair of double-strand breaks (DSBR) by non-homologous end joining (NHEJ) (22, 45, 53). We and others (54) do not find homologs for the NHEJ proteins, KU70, KU80 and DNA ligase IV in *P. falciparum* and *T. parva* and *B. bovis* only contains DNA ligase IV. Indeed, these organisms are known to predominantly employ homologous recombination for DNA repair (21). The NHEJ proteins are present in *T. gondii* and the other coccidians and we have validated their role in NUOT insertion in *T. gondii* by examining repair of a CRISPR induced DSB in a Δ*ku80* strain. This strain was created because *T. gondii* prefers NHEJ over other methods for DSB repair (48). The phylogenetic profile of the NHEJ pathway genes and the fact that *T. gondii* uses NHEJ preferentially suggests that the proficiency of the NHEJ pathway is sufficient to explain the high NUOT content in *T. gondii*. However, it doesn’t explain the variation in insertion levels observed between coccidian species.

We have shown that the NUMT content even within closely related apicomplexan species varies significantly. These data are largely consistent with previous observations where a large variation in NUMT content has been described among *Drosophila melanogaster*, *Anopheles gambiae*, and *A. mellifera*, and even among mammals like human, mouse and rat (16, 17, 41, 55-57). This variation may be explained by two major forces: differences in the frequency at which species acquire and retain DNA from their mitochondria and the differential rates of NUMT removal (56, 58). The frequency at which mtDNA is transferred to the nucleus can be influenced by many factors, including the total number of mitochondria within a given cell, and the level of vulnerability to stressful agents that may damage the mitochondria thereby releasing mtDNA. However, given that apicomplexans generally contain only one mitochondrion (and apicoplast) per cell, this cannot sufficiently explain the observable differences in NUMT (and NUPT) accumulation (59). There is one significant difference between *Eimeria* and *Sarcocystis* which have fewer NUOTs and the high NUOT group of *T. gondii, H. hammondi* and *N. caninum,* the former species contain transposable elements (29, 60) whereas the others do not. Perhaps *Sarcocystis* and *Eimeria* have evolved mechanisms for keeping the insertion of DNA under control and the *Toxoplasmatinae* lost this ability when they cleared mobile elements from their genome. Alternatively, *the Toxoplasmatinae* may benefit from this high level of genome variation precisely because they have lost their mobile elements and the sources of genetic variation they provide. *Toxoplasma* in particular, also has the added biology of being able to propagate asexually. This means it can spend long periods of time as a haploid organism without a homologous chromosome to use as a repair template, which could create additional advantages to having a steady supply of organellar DNA patches. There are many hypotheses that deserve exploration.

A major unanswered question is how organellar DNA enters the nucleus and is available for NHEJ repair. Analysis of existing NUOTs suggests that smaller fragments (100-200 bp) are most frequently observed, but that significantly longer inserts are occasionally seen. NUMT inserts appear to be random genome segments and do not correspond to the unique SBs of the *T. gondii* mtDNA. It is possible that oxidative stress in the mitochondria may play a role in the generation of random mtDNA fragments as single and double strand mtDNA breaks are observed in yeast (61–63). But this explanation will not work for ptDNA fragments. The CRISPR DSB repair experiment revealed slight differences in the insert lengths depending on insert DNA source with CRISPR plasmid DNA inserts being longer than ptDNA which were slightly longer than mtDNA inserts. In this experimental setting, NUPTs were also observed slightly more frequently than NUMTs a finding that is at odds with the much higher abundance of NUMTs in *T. gondii* and all the other coccidian genomes. The reason(s) why NUMTs greatly exceed NUPTs is unclear but likely reflects cell biological differences in these independently acquired organelles. There are observed differences in the calculated ages of NUOTs in the *Toxoplasmatinae* suggesting that different lineages have experienced different waves of insertion or variation in removal that could not be recapitulated in the experiments used here. We did observe the insertion of multiple non-continuous DNA fragments including DNA from different sources both at the induced CRISPR DSB and naturally occurring in the genome sequence attesting to the efficiency of DSB repair in *T. gondii*. These mosaic inserts, termed NUMINs have been observed previously in plants (64).

Our attempt to identify NUOTs that were not ubiquitous in the 16 *T. gondii* strains examined revealed 50 insertions in or near genes that were enriched in a Gene Ontology analysis for processes related to microtubules and nucleotides. The distribution of NUMTs among the strains revealed both recent insertion and patterns that are most easily explained as the result of sexual recombination. Our search for orthologous NUOTs between *T. gondii* and *N. caninum* resulted in only 5 orthologous NUMTs and no orthologous NUPTs. This finding highlights the efficiency with which NUOTs are removed from the genome. These species diverged an estimated 28 MY ago (43). Of the 5 detectable NUMTs, one was more conserved relative to surrounding sequence, suggesting it may have evolved under functional constraint and might shed light on the types of evolutionary innovation NUOTs can provide. Indeed, functional assays suggest it might act as a cis-regulatory element affecting promoter activity of the U6 snRNA associated Sm family protein as promoter activity was reduced upon NUMT removal in a reporter construct. All coccidians have a large number of introns thus, one could speculate that this might confer an effect, but this would need to be tested. Surprisingly, the *T. gondii*-specific NUMT which is inserted near the putative myosin heavy chain ATPase gene in all strains examined dramatically decreases the gene promoter activity. These findings are intriguing because they suggest that both ancient and recent NUMT insertions can influence gene regulation. Because the putative myosin heavy chain ATPase gene implicated here belongs to a diverse family of proteins involved in parasite motility, division and penetration of host cells (65), it is possible that the NUMT inserted within its promoter region, could modulate the pathogenicity and virulence of the parasite, but this remains to be proven. NUMTs are increasingly associated with functional consequences, particularly in human diseases (52) and mitigating deletions in the evolution of primates (45). For example, mtDNA instability and increased NUMT somatic integration has been associated with colorectal adenocarcinoma (66).

## Conclusion

Following identification of both *T. gondii* organellar genome sequences, it became possible to characterize the NUMTs and NUPTs that have hindered some molecular studies of these organellar DNAs. Shockingly, this protist pathogen and its closest relatives *H. hammondi* and *N. caninum* harbor the highest NUOT density ever observed. While most NUOTs exhibit various levels of sequence decay, several inserts are identical in sequence to their organellar source suggesting they are recent and that nuclear insertion and turnover of organellar DNA is an ongoing process in these species. NHEJ was shown to be essential for repair of an induced DSB. Examination of 16 *T. gondii* strains revealed 56 insertions that are differentially present and a search for orthologous insertions in *N. caninum* yielded 5 NUMTs. Deletion of one orthologous and one non-orthologous NUMT affected gene expression of reporter constructs. To our knowledge this is the first study to experimentally establish that NUMTs can also influence gene expression. NUOTs provide a previously unexplored source of genomic variation in these important pathogens.

## Materials and Methods

### Source of genome Sequences

The nuclear genomic sequences of *Toxoplasma gondi* ME49 (NCBI assembly GCA_000006565.1)*, Hammondia hammondi* H.H.34 (GCA_000258005.1) *Neospora caninum* Liverpool (GCA_000208865.1), *Sarcocystis neurona* SN3 (GCA_000727475.1)*, and Eimeria tenella* Houghton (GCA_000499545.1) were downloaded from ToxoDB.org (44). *Plasmodium falciparum* 3D7 (GCA_000002765.1)*, Babesia bovis* (GCA_000165395.1) and *Theileria annulata* (GCA_000003225.1) nuclear as well as organellar genome sequences were obtained from PlasmoDB.org and PiroplasmaDB.org respectively.

### Identification of NUMTs and NUPTs

RepeatMasker version 4.0.5 (http://www.repeatmasker.org) was used to identify organellar DNA sequences in the nuclear genome sequences. mtDNA and ptDNA seqeunces were used as the “repeat library” and “cross_match” was used with filtering out low complexity sequences and other parameters as default, to mask the corresponding nuclear genomes to identify NUMTs and NUPTs respectively. For *T. gondii* and *N. caninum* the species-specific 21 sequence blocks as well as cytochrome coding sequences, *coxI*, *coxIII* and *cob* (36) were used to mask the nuclear chromosomal sequences. *T. gondii* apicoplast sequence (GenBank: U87145.2) was used to detect NUPTs in *T. gondii* ME49 and *N. caninum* Liverpool. *T. gondii* mtDNA and ptDNA sequences were used to identify NUOTs in *H. hammondi*. The partial cytochrome sequences annotated in *S. neurona* and *T. gondii* mtDNA sequences were used to identify NUMTs in *S. neurona* and the *S. neurona* ptDNA sequence was used to identify NUPTs. NUMTs and NUPTs in *E. tenella* were detected using its own organellar sequences (GenBank Accessions: AB564272.1 and AY217738.1. Similarly, *Plasmodium, Babesia* and *Theileria* species-specific organellar sequences obtained from VEuPathDB.org were used to screen their respective nuclear chromosome sequences for NUOTs and genes involved in the NHEJ DSB repair pathway. NUOTs shorter than 28 bp and overlapping insertions were only counted once.

### Identification of NUMT location

Coordinates of coding regions, introns, exons, intergenic and 1 kb upstream and downstream flanking regions were obtained from the annotation data available on ToxoDB.org release 13 (44). Based on coordinates, a custom script was used to identify the genomic location for each NUMT. NUMTs located in non-coding exons were classified as being present in the UTR. NUMTs spanning more than one genomic feature were classified separately. Insertion sites were inspected to see if a target site preference could be observed.

### Identification of segmental duplications, shared and uniquely-present NUMTs

Segmental duplications were identified by extracting NUMT sequences along with 150 bp of the flanking regions. The extracted sequences were used as BLASTN queries against themselves. If a NUMT along with at least 100bp of flanking sequence was present at least twice it was counted as a segmental duplication (27). Following the identification NUMTs in *T. gondii* ME49 reference nuclear genome, NUMTs were similarly identified in nuclear genomes of the fully assembled *T. gondii* strains, GT1 and VEG and *de novo* assembled sequences of ARI, CAST, TgCtBr5, TgCtBr9, TgCtPRC2, COUG, TgCtCo5, FOU, GAB2-2007-GAL-DOM2, MAS, p89, RUB and VAND, obtained from ToxoDB.org (44). Any scaffold or contig that contained only mtDNA sequence was excluded. In order to identify NUMTs that were shared or unique to a strain, each NUMT sequence along with 200 bp of flanking sequences was searched against the NUMTs with 200 bp flanking sequences of all other strains using BLASTN. NUMTs that were present in only one strain were called strain-specific *T. gondii* NUMTs. NUMTs that are present in all are called ‘shared’. Strain-specific NUMTs in ME49, GT1 and VEG were manually verified. A similar approach was utilized to identify NUMTs that were shared or uniquely present among *T. gondii* ME49 and *N. caninum*. All BLASTN search results were filtered for an E-value = 0 and query coverage of >90%.

### Estimation of NUOT age

The relative date of insertion of a NUMT can be inferred from the NUMTs age. RepeatMasker reports a percent divergence between the NUMT and the corresponding mtDNA sequence. The percent divergences were converted into genetic distances using the Jukes-Cantor Model formula

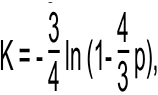

where K is the genetic distance and p is the percent divergence between the mtDNA and NUMT. Percent divergence was calculated by dividing the number of substitutions between the two sequences by the sequence length. Using the genetic distance the age of each NUMT was calculated using the formula, T = K/2μ, where T is the age of the NUMT, K is the genetic distance (Dataset S1), μ is the intron mutation rate. The intron mutation rate was used to calculate the age of the NUMTs since NUMTs are not expected to be under selective pressure and should have a neutral mutation rate similar to the introns. The intron mutation rate was previously calculated as 2.12 × 10^-8^ (10). The same rate was for used for all three species.

### Expression constructs and Transient Transfection

PCR primers (Table S6) containing the attB1 and attB2 sites were used to amplify the appropriate promoter and 5′-UTR regions from ME49 genomic DNA for the two genes tested. A two-step overlap-extension PCR technique (67) was employed to delete the NUMT from each promoter. Following sequencing verification, these promoter fragments were moved into a Firefly luciferase-expressing vector. As constitutive *T. gondii* promoter (α-tubulin)-driven Renilla luciferase-expressing construct was co-transfected along with the experimental construct. Parasite culture and transient transfections were performed as described (68). A dual luciferase assay was performed on the extract using the Promega Dual Luciferase kit. Reporter activity from the WT or ΔNUMT promoter was measured relative to the internal control. Each electroporation experiment was performed 6 times and luciferase assays were performed in duplicate for expression measurements. The unpaired Students *t*-test was used to calculate the statistically significant difference in expression levels between WT and mutagenized promoter activity; *p* < 0.05 was considered statistically significant.

### CRISPR-Cas9 transfection assays, amplicon sequencing and analysis of inserts

*T. gondii* strains RH and an RH mutant TATiΔ*ku80*, referred here as WT and Δ*ku80* (69) respectively were cultured on human immortalized fibroblast cells grown in Dulbecco’s Modified Eagle’s Medium supplemented with 10% heat-inactivated Cosmic Calf Serum, 0.5% 10mg/ml penicillin-streptomycin and 0.05% 10 mg/ml gentamycin. *T. gondii* RH WT and Δ*ku80* parasites were filtered, washed and resuspended in cytomix (supplemented with ATP and glutamine) at a concentration of 3.3 ×10^7^ parasites/ml. 10^7^ parasites were electroporated with 8 μg of the CRISPR/Cas9 plasmid with a single guide RNA targeting the UPRT locus utilizing a BTX electroporator. hTERT cells were infected with the transfected parasites (WT or Δ*ku80* in triplicate) and grown for 48 hr without drug selection. After 48 hr, lysed parasites were passed to fresh hTERT cells and placed under 10 μM of 5-fluorodeoxyuridine (5-FUDR) drug selection to select for parasites with a non-functional UPRT gene. Parasites were harvested following four rounds of drug selection and DNA was extracted using Qiagen Blood and Tissue kit. PCR amplicons of the UPRT locus from DNA extracted from all six transfections were generated. Amplicons were indexed and Illumina MiSeq PE-150 bp sequencing was performed. Filtered and trimmed paired-end reads were then merged using FLASH (default parameters) (70, 71). Unique reads were identified using the usearch fastx_uniques script (72). The unique reads were then investigated for the presence of *T. gondii* mtDNA (21 sequence blocks), ptDNA, and CRISPR/Cas9 plasmid DNA sequences using RepeatMasker and each genome sequence as a repeat library. RepeatMasker results were then analyzed to identify the characteristics of the inserts reported in Fig. 4.

## Acknowledgments

We thank Drs. Michael Cipriano (University of Georgia) and Boris Striepen (University of Pennsylvania) for generously providing the RH mutant TATiΔ*ku80* and the CRISPR/Cas9 plasmid. This work was funded by NIH R01AI068908 to JCK.

## Data sharing

Amplicon sequences are available via NCBI BioProject PRJNA961209.

## Supplementary Information Text

### Supplementary Materials and Methods

#### DNA extraction and PCR

DNA from *T. gondii* RH and ME49 parasites was extracted using DNeasy Blood and Tissue Kit (Qiagen) following manufacturer’s cultured cells protocol. PCR was performed using the following conditions using primers in Table S6: initial denaturation at 95°C for 3 minutes; followed by 35 cycles of amplification at 95°C for 60 seconds, 52.5°C for 45 seconds, and 72°C for 40 seconds. The PCR products were visualized using agarose gel electrophoresis.

#### Identification of NUMTs and NUPTs

RepeatMasker version 4.0.5 (http://www.repeatmasker.org) rather than the frequently used BLASTN was used to identify organellar DNA sequences in the nuclear genomes as it provides better accuracy, handles many-to-one hits, avoids overestimates and provides a detailed annotation of the identified sequences, which facilitates subsequent characterization. Mitochondrial and apicoplast genome sequences were used as the “repeat library” and “cross_match” was used as the search engine, with filtering out low complexity sequences and other parameters as default, to mask the corresponding nuclear genomes to identify NUMTs and NUPTs respectively. For *T. gondii* and *N. caninum* the species-specific 21 sequence blocks as well as cytochrome coding sequences, *coxI*, *coxIII* and *cob* (1) were used to mask the corresponding assembled nuclear chromosomal sequences. Since the 21 sequence blocks occur in a number of random arrangements and a NUMT can arise from a non-contiguous mitochondrial sequence, the cytochrome sequences were included to minimize the overestimation of the number of NUMTs. The apicoplast sequence *T. gondii* (GenBank: U87145.2) was used to detect NUPTs in *T. gondii* ME49 and *N. caninum* Liverpool. Since the mtDNA sequence could not be distinguished from NUMTs in *H. hammondi* (ToxoDB.org) (2), *T. gondii* mtDNA sequences were used to identify NUMTs in *H. hammondi*. Additionally, the publicly available apicoplast sequence for *H. hammondi* is incomplete in comparison to the well annotated plastid sequence of *T. gondii.* Therefore, *T. gondii’s* apicoplast sequence was used to screen for NUPTs in *H. hammondi*. The partial cytochrome sequences annotated in *S. neurona* and *T. gondii* mtDNA sequences were used to identify NUMTs in *S. neurona* and the parasites’ own apicoplast DNA sequence was used to identify NUPTs. NUMTs and NUPTs in *E. tenella* were detected using its own organellar genome sequences (GenBank Accessions: AB564272.1 and AY217738.1. Similarly, *Plasmodium, Babesia* and *Theileria* species-specific organellar sequences obtained from VEuPathDB.org were used to screen their respective nuclear chromosome sequences for NUOTs and genes involved in the NHEJ DSB repair pathway. NUOTs shorter than 28 bp and overlapping insertions were only counted once.

#### Identification of segmental duplications, shared and uniquely-present NUMTs

Segmental duplications were identified by extracting NUMT sequences along with 150 bp of the flanking regions. The extracted sequences were used as BLASTN queries against themselves. If a NUMT along with at least 100bp of flanking sequence was present at least twice it was counted as a segmental duplication (3).

Following the identification NUMTs in *T. gondii* ME49 reference nuclear genome, NUMTs were similarly identified in nuclear genomes of two other fully assembled *T. gondii* strains, GT1 and VEG using the same mtDNA library repeat described above. Subsequently, the *de novo* assembled genomes of 13 *T. gondii* strains namely, ARI, CAST, TgCtBr5, TgCtBr9, TgCtPRC2, COUG, TgCtCo5, FOU, GAB2-2007-GAL-DOM2, MAS, p89, RUB and VAND, were downloaded from ToxoDB.org (2) and NUMTs were similarly identified. It should be noted that the genomes of these 13 strains have not be assembled into chromosomes and occur as scaffolds and contigs. Therefore, NUMTs that are present at the ends of these sequences could not be included in the analyses due to the lack of flanking regions. Additionally, any scaffold or contig that contained only mitochondrial DNA sequence was not included in further analysis. In order to identify NUMTs that were shared or unique among the strains, each NUMT sequence along with 200 bp of flanking sequences from one strain was searched against the NUMTs with 200 bp flanking sequences of all other strains using BLASTN. For each pairwise comparison, the NUMT was considered to be shared only when >150 bp of the upstream and downstream flanking sequence was present. If both flanking sequences were not present, the match was not considered. NUMTs that were present in only one strain were called ‘strain-specific *T. gondii* NUMTs’ and NUMTs that present in more than one strain were called ‘differentially present *T. gondii* NUMTs’. NUMTs that were present in all 16 strains were called ‘shared *T. gondii* NUMTs’. Strain-specific NUMTs identified from the comparison of just the three fully-assembled strains ME49, GT1 and VEG were manually verified. Finally, a similar approach was utilized to identify NUMTs that were shared or uniquely present among *T. gondii* ME49 and *N. caninum*. All BLASTN search results were filtered for an E-value = 0 and query coverage of >90%.

#### Estimation of NUOT age

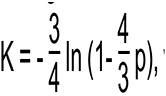

where K is the genetic distance and p is the percent divergence between the mtDNA and NUMT. Percent divergence was calculated by dividing the number of substitutions between the two sequences by the sequence length. Using the genetic distance the age of each NUMT was calculated using the formula, T = K/2μ, where T is the age of the NUMT, K is the genetic distance (Dataset S1), μ is the intron mutation rate. The intron mutation rate was used to calculate the age of the NUMTs since NUMTs are not expected to be under selective pressure and should have a neutral mutation rate similar to the introns. The intron mutation rate was previously calculated as 2.12 × 10^-8^ (4). The same rate was for used for all three species.

#### Expression constructs and Transient Transfection

PCR primers (Table S6) containing the attB1 and attB2 sites were used to amplify the appropriate promoter and 5′-UTR regions from ME49 genomic DNA for the two genes tested. A two-step overlap-extension PCR technique (5) was employed to delete the NUMT sequence from each promoter. The Gateway™ cloning system (Invitrogen) was used to clone the WT and deletion promoters. These two kinds of promoters were first cloned into pDONR221 via the BP reaction. Following sequencing verification, these promoter fragments were moved into a firefly luciferase-expressing vector (destination vectors) via the LR reaction. As constitutive *T. gondii* promoter (α-tubulin)-driven renilla luciferase-expressing construct was co-transfected along with the experimental construct. Nucleotide positions in these deletion studies are referenced with respect to the start of translation (+1). Parasite culture and transient transfections were performed as described (6). In short, *T. gondii* RH tachyzoites were transiently transfected via electroporation. 18-24 hours post-electroporation the cells were scraped and lysed using passive lysis buffer (Promega, Madison, WI, USA) and a dual luciferase assay was performed on the extract using the Promega Dual Luciferase kit. The different substrate requirements for Firefly and Renilla luciferase allowed us to assay reporter expression for each construct sequentially within the same extract. Reporter activity from the WT or mutagenized promoter was measured relative to the internal control, eliminating errors due to variation in parasite populations and individual transfections. Each electroporation experiment was performed 6 times and luciferase assays were performed in duplicate for expression measurements. The unpaired Students *t*-test was used to calculate the statistically significant difference in expression levels between WT and mutagenized promoter activity; *p* < 0.05 was considered statistically significant.

#### CRISPR-Cas9 transfection assays, amplicon sequencing and analysis of inserts

*T. gondii* strains RH and an RH mutant TATiΔ*ku80*, referred here as WT and Δ*ku80* (7) respectively were cultured on human immortalized fibroblast (hTERT) cells grown in Dulbecco’s Modified Eagle’s Medium (DMEM) supplemented with 10% heat-inactivated Cosmic Calf Serum, 0.5% 10mg/ml penicillin-streptomycin and 0.05% 10 mg/ml gentamycin. *T. gondii* RH WT and Δ*ku80* parasites were filtered, washed and resuspended in cytomix (supplemented with ATP and glutamine) at a concentration of 3.3 ×10^7^ parasites/ml. 10^7^ parasites were electroporated with 8 μg of the CRISPR/Cas9 plasmid with a single guide RNA targeting the UPRT locus utilizing a BTX electroporator (CRISPR/Cas9 plasmid and guide RNA kindly provided by Michael Cipriano and Boris Striepen). hTERT cells were infected with the transfected parasites (WT or Δ*ku80* in triplicate) and grown for 48 hours without drug selection. After 48 hours, lysed parasites were passed to infect fresh hTERT cells and placed under 10 μM of 5-fluorodeoxyuridine (5-FUDR) drug selection to select for parasites with a mutation in the UPRT gene. Parasites were harvested following four rounds of drug selection and DNA was extracted using Qiagen Blood and Tissue kit following the manufacturer’s protocol. PCR was performed using primers (Forward primer: 5′-TATGGTAATTGTGAAGTGACAACCCCTCTGGA-3′; Reverse primer: 5′-AGTCAGTCAGCCCAAGCCACTTTCCATCGACT-3′) targeting the UPRT locus to generate 66 bp amplicons from DNA extracted from all six transfections. Amplicons were then indexed and paired-end sequencing (PE 150 bp) was performed using the Illumina MiSeq sequencer.

Sequenced paired-end reads were filtered and trimmed using trimmomatic (minimum length:36, sliding window:4:20) (8), The processed paired-end reads were then merged using FLASH (default parameters) (8, 9). Following quality control and merging of paired-end reads unique reads were identified using the usearch fastx_uniques script (10). The unique reads were then investigated for the presence of *T. gondii* mtDNA (21 sequence blocks), ptDNA, and CRISPR/Cas9 plasmid DNA sequences using RepeatMasker and each genome sequence as a repeat library. RepeatMasker results were then analyzed to identify the characteristics of the inserts reported in Fig. 4.

---

**Figure S1.**
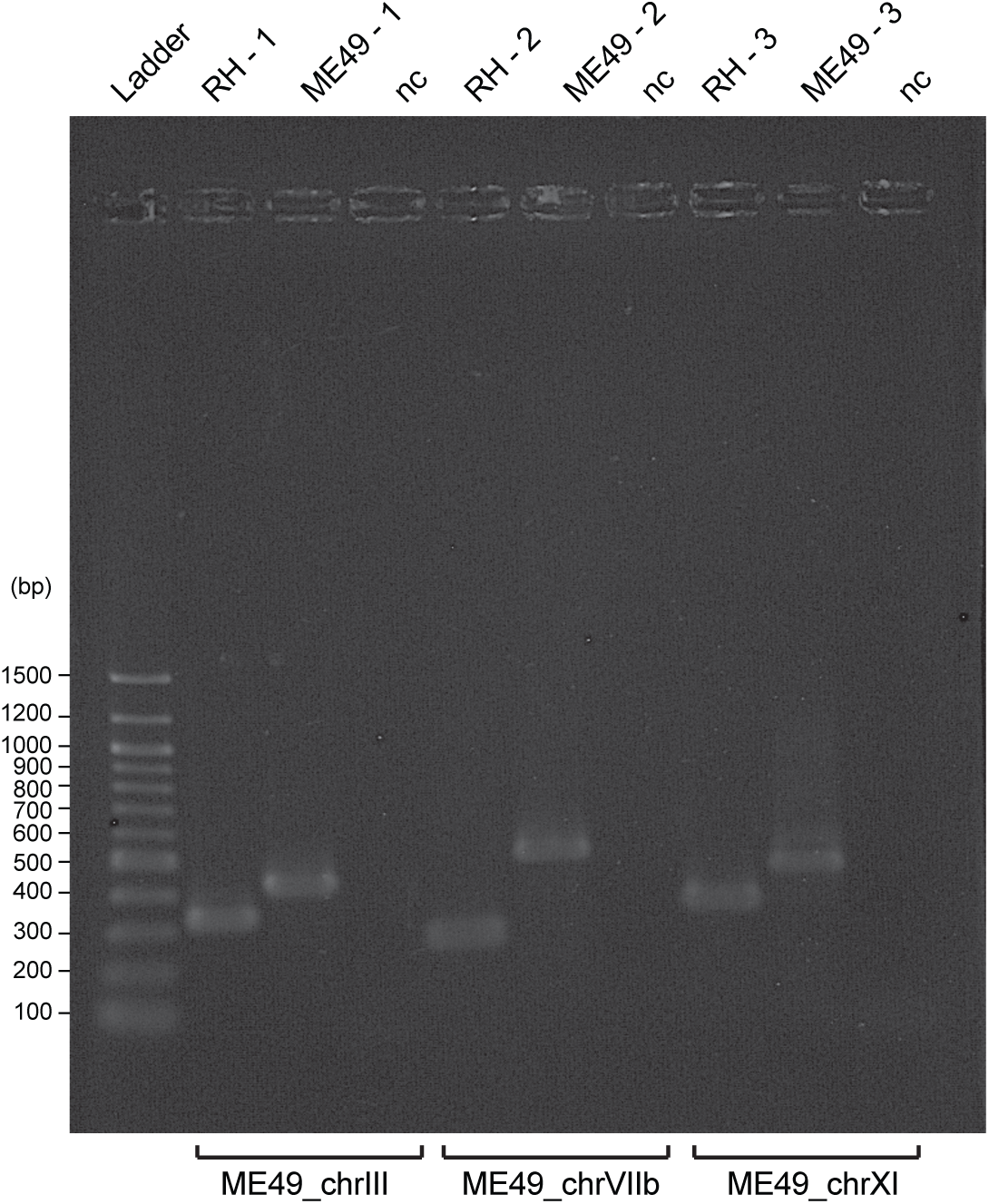
PCR verification of select strain-specific NUMTs. Genomic DNA from *T. gondii* strains ME49 and RH were assayed with three primers sets that each target a strain specific NUMT loci. The strain is indicated across the top and ‘nc’ denotes the negative control for each reaction. The primers are specified on the bottom and the primer sequences are provided Table S6. ME49_chrIII ME49_chrVIIb and ME49_chrXI correspond to NUMTs located at TGME49_chrIII:1730617..1730859, TGME49_chrV11b:1710367..1710510, and TGME49_chrXI:3550511..3550993 in Table S4 respectively. All NUMTs shown here are present in ME49 and absent in RH. RH was utilized instead of GT1 since RH is a widely used *T. gondii* strain in laboratory cultures. RH and GT1 are closely related strains. However, a fully assembled genome sequence is unavailable.

**Figure S2.**
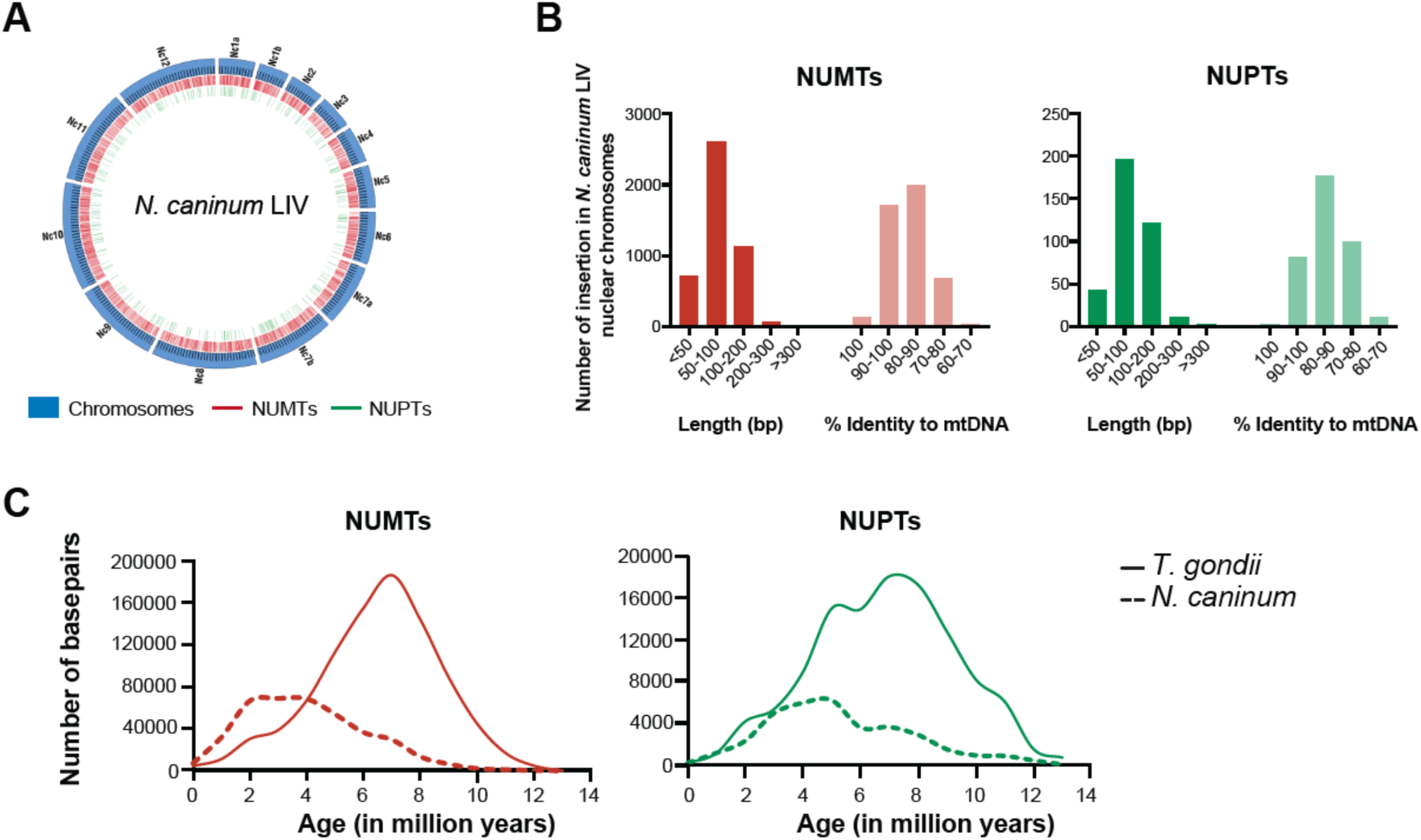
Characteristics of NUMTs and NUPTs in *N. caninum LIV*. *(A)* Circos plot represents the distribution of NUMTs and NUPTs in the nuclear chromosomes of *N. caninum LIV*. The blue bands indicate the chromosomes as labeled and each tick on the band equals 100 kb. The ticks interior to the chromosomes are indicate the location of NUMTs and NUPTs as referenced in the key. *(B)* The length and the percent identity of a NUMT and NUPT to *N. caninum* mt and ptDNA respectively was calculated from RepeatMasker results and the distribution is plotted. *(C)* The age of each NUMT and NUPT was calculated based on the percent divergence from the mt and ptDNA as described in the Methods and the distribution of the age is plotted against the number of base pairs.

**Figure S3.**
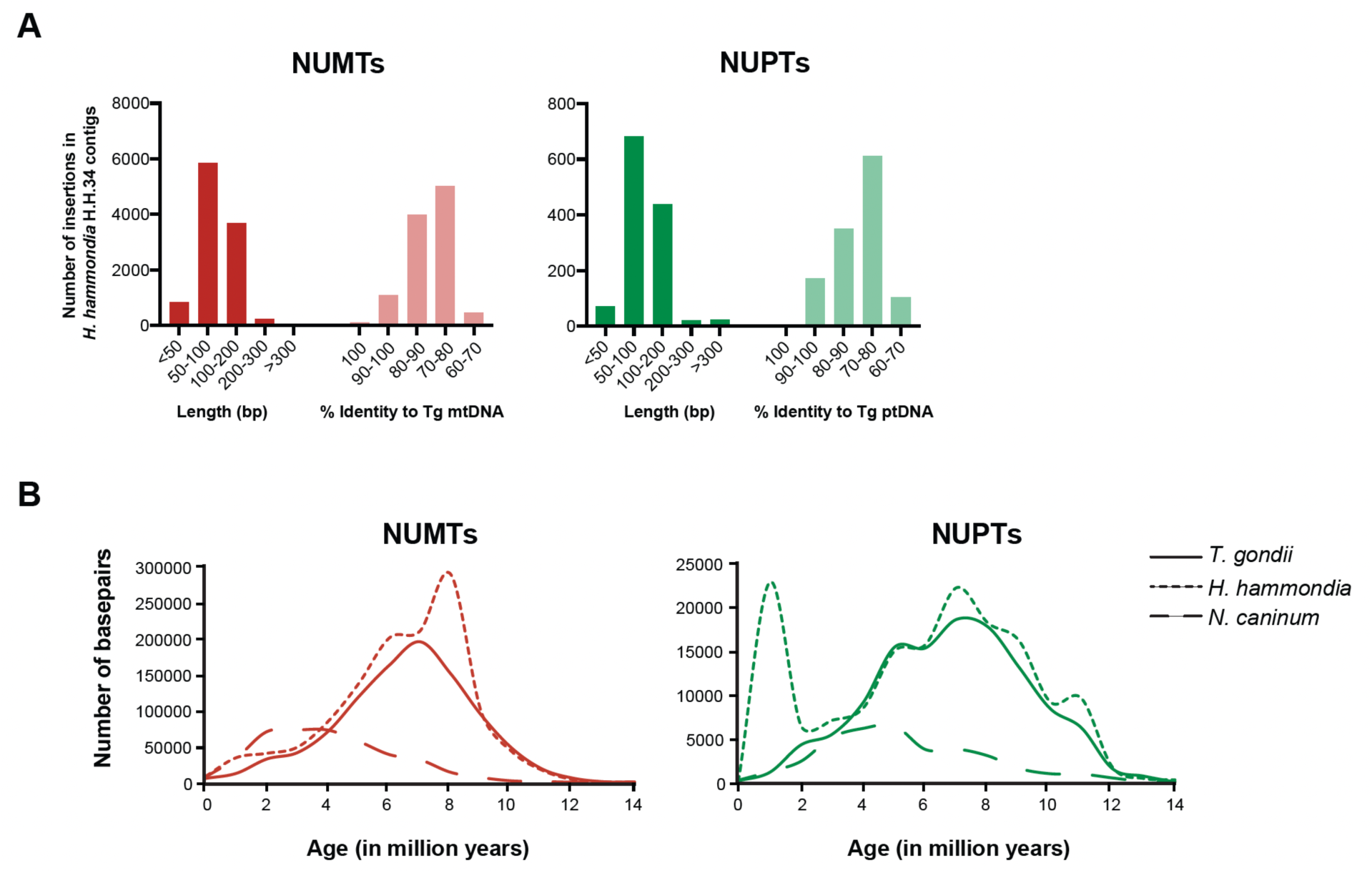
Characteristics of NUMTs and NUPTs in *H. hammondi* H.H.34. (A) The length and percent identity of a NUMT and NUPT to *T. gondii* mt and ptDNA respectively was calculated from RepeatMasker results and the distribution is plotted. (B) The age of each NUMT and NUPT was calculated based on the percent divergence from the mt and ptDNA as described in the Methods and the distribution of the age is plotted against the number of base pairs. It should be noted that the NUM/PT analysis in *H. hammondi* was performed on genome contigs as a chromosome assembly is unavailable. Thus, the spike in the number of NUPT base pairs at ∼1 million years is likely due to the presence of unassembled apicoplast genome sequences.

**Figure S4.**
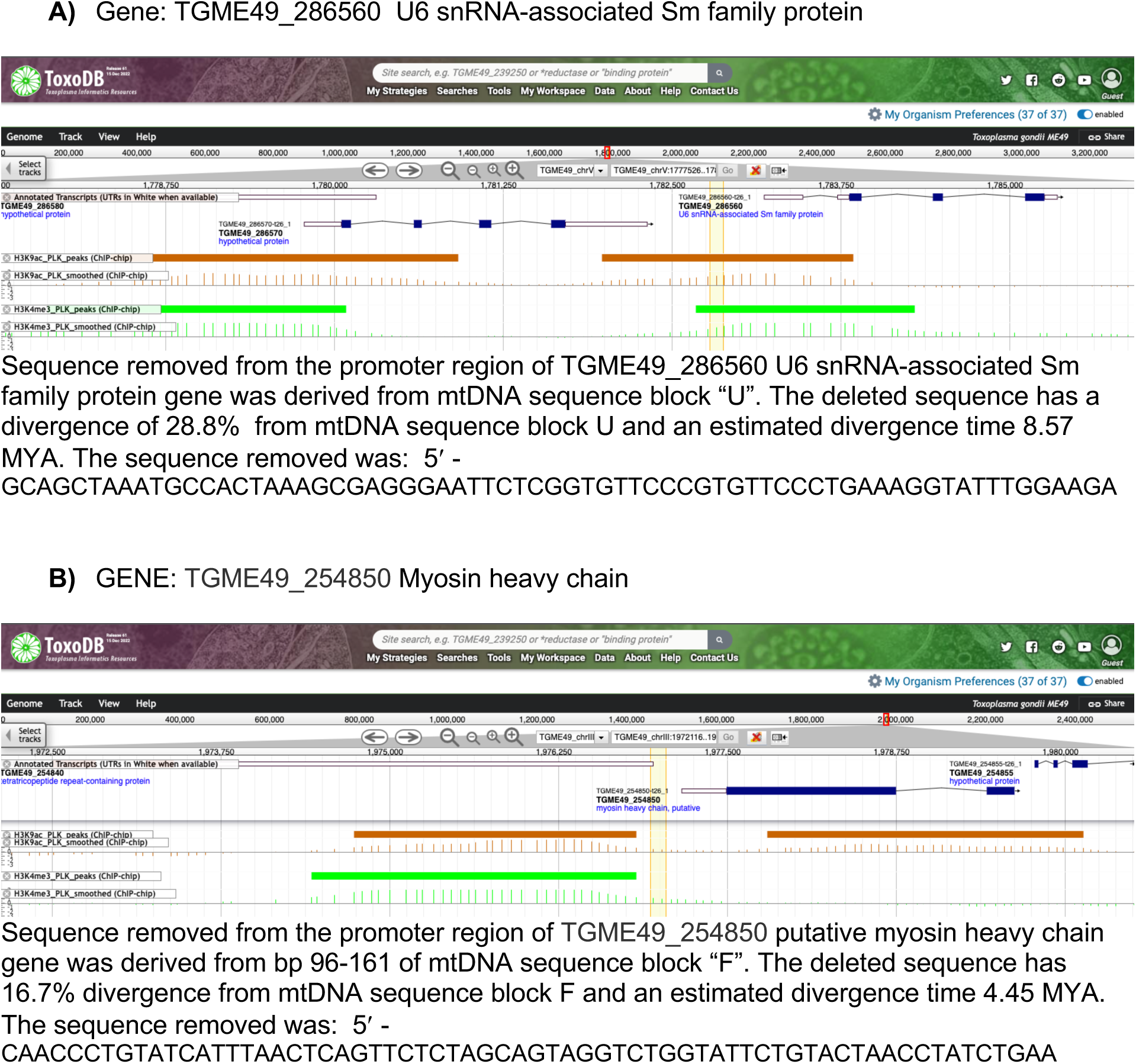
NUMT location and epigenetic marks for promoters used in transient *in vitro* reporter expression assays. (A) Promoter region of TGME49_286560 U6 snRNA-associated Sm family protein gene that shares an orthologous conserved NUMT derived from mtDNA sequence block “U” with *N. caninum*. (B) Promoter region of TGME49_254850 Myosin heavy chain gene that is not orthologous and was derived from mtDNA sequence block “F”. Screen shots are from ToxoDB.org. Yellow highlight indicates the location of the NUMT. Orange and green data tracks represent H3K9ac and H3K4Me3 ChIP-chip data respectively. Both genes are located on the forward strand. The CDS is colored solid blue in the gene models. The uncolored regions on the ends of the gene models represent the 5′ and 3′ UTRs.

**Table S1.** Annotation of the 3,369 bp strain-specific NUMT on ChrX in *T.g*. ME49

| **Chromosome** | **Start** | **End** | **Divergence**<br>**from**<br>**mtDNA (%)** | **mtDNA**<br>**gene/block** | **Start** | **End** |
| --- | --- | --- | --- | --- | --- | --- |
| TGME49_chrX | 5732558 | 5732660 | 0 | U | 1 | 103 |
| TGME49_chrX | 5732662 | 5732907 | 1.6 | coxIII | 216 | 459 |
| TGME49_chrX | 5732891 | 5732976 | 0 | O | 1 | 86 |
| TGME49_chrX | 5732977 | 5733228 | 0 | K | 1 | 252 |
| TGME49_chrX | 5733227 | 5733447 | 0 | cob | 1 | 221 |
| TGME49_chrX | 5733451 | 5733767 | 0.3 | coxIII | 1 | 317 |
| TGME49_chrX | 5733697 | 5734062 | 0 | coxI | 11 | 376 |
| TGME49_chrX | 5734077 | 5734453 | 0.5 | cob | 20 | 396 |
| TGME49_chrX | 5734448 | 5734516 | 0 | H | 1 | 69 |
| TGME49_chrX | 5734467 | 5734581 | 0 | coxI | 1377 | 1491 |
| TGME49_chrX | 5734580 | 5734683 | 4.9 | coxIII | 216 | 317 |
| TGME49_chrX | 5734582 | 5734666 | 0 | J | 1 | 85 |
| TGME49_chrX | 5734667 | 5734752 | 0 | O | 1 | 86 |
| TGME49_chrX | 5734753 | 5734931 | 0 | F | 1 | 179 |
| TGME49_chrX | 5734932 | 5735016 | 0 | R | 1 | 85 |
| TGME49_chrX | 5735033 | 5735098 | 0 | U | 1 | 66 |
| TGME49_chrX | 5735100 | 5735284 | 0 | coxIII | 275 | 459 |
| TGME49_chrX | 5735285 | 5735354 | 0 | M | 457 | 526 |
| TGME49_chrX | 5735351 | 5735396 | 0 | D | 1 | 46 |
| TGME49_chrX | 5735397 | 5735508 | 0 | K | 334 | 445 |
| TGME49_chrX | 5735509 | 5735860 | 0 | U | 1 | 353 |
| TGME49_chrX | 5735862 | 5735926 | 0 | coxIII | 395 | 459 |

**Table S2.** Characteristics of the NUMTs generated from the 21 mtDNA blocks in *T. gondii* and *N. caninum*

| Block | *T. gondii* ME49<sup>a</sup> |  |  |  |  |  |  |  | *N. caninum* LIV<sup>a</sup> |  |  |  |  |  |  |  | Entire block found at 100% identity in nuclear genome? |  |
| --- | --- | --- | --- | --- | --- | --- | --- | --- | --- | --- | --- | --- | --- | --- | --- | --- | --- | --- |
|  | Length of block | # of NUMTs | # of bp | % of genome | Number of insertions based on % identity to mtDNA<sup>b</sup> |  |  |  | Length of block | # of NUMTs | # of bp | % of genome | Number of insertions based on identity to mtDNA<sup>b</sup> |  |  |  |  |  |
|  |  |  |  |  | 100 | >=90 | >=80 | >=70 |  |  |  |  | 100 | >=90 | >=80 | >=70 | *Tg* | *Nc* |
| A | 196 | 298 | 24376 | 0.04 | 1 | 42 | 158 | 293 | 196 | 161 | 12205 | 0.02 | 6 | 65 | 136 | 161 | N | N |
| B | 40 | 31 | 1213 | 0.002 | 4 | 15 | 31 | 31 | 40 | 39 | 1538 | 0.002 | 6 | 31 | 39 | 39 | Y | Y |
| C | 1050 | 246 | 28107 | 0.04 | 2 | 26 | 84 | 223 | 1050 | 361 | 31440 | 0.05 | 6 | 110 | 291 | 357 | N | N |
| D | 82 | 181 | 11544 | 0.02 | 4 | 25 | 115 | 178 | 82 | 78 | 4710 | 0.008 | 3 | 48 | 75 | 78 | N | N |
| E | 683 | 829 | 88673 | 0.14 | 5 | 64 | 345 | 783 | 683 | 414 | 38823 | 0.07 | 16 | 168 | 333 | 410 | N | N |
| F | 179 | 308 | 25785 | 0.04 | 2 | 35 | 166 | 297 | 179 | 188 | 14559 | 0.03 | 4 | 74 | 163 | 188 | Y | N |
| H | 447 | 291 | 28704 | 0.05 | 1 | 36 | 123 | 274 | 447 | 162 | 13416 | 0.02 | 4 | 58 | 141 | 162 | N | N |
| I | 204 | 160 | 13483 | 0.02 | 1 | 21 | 92 | 152 | 204 | 71 | 5274 | 0.009 | 0 | 36 | 61 | 71 | N | N |
| J | 85 | 626 | 42485 | 0.07 | 5 | 68 | 385 | 624 | 85 | 303 | 18362 | 0.03 | 19 | 168 | 283 | 303 | Y | Y |
| K | 445 | 696 | 64432 | 0.1 | 6 | 69 | 334 | 668 | 445 | 292 | 24364 | 0.04 | 12 | 138 | 248 | 288 | N | N |
| L | 159 | 610 | 53125 | 0.08 | 3 | 45 | 288 | 588 | 159 | 278 | 21453 | 0.04 | 13 | 122 | 233 | 277 | Y | Y |
| M | 754 | 890 | 86617 | 0.14 | 9 | 87 | 393 | 836 | 754 | 406 | 37383 | 0.06 | 11 | 163 | 336 | 403 | N | N |
| N | 166 | 173 | 15536 | 0.02 | 0 | 14 | 78 | 170 | 166 | 123 | 9706 | 0.02 | 3 | 42 | 103 | 123 | N | N |
| O | 86 | 234 | 15987 | 0.03 | 3 | 33 | 137 | 234 | 86 | 130 | 8091 | 0.01 | 6 | 50 | 122 | 130 | Y | Y |
| P | 184 | 301 | 26339 | 0.04 | 0 | 43 | 159 | 281 | 184 | 146 | 11253 | 0.02 | 6 | 58 | 126 | 146 | N | N |
| Q | 65 | 62 | 3489 | 0.01 | 2 | 15 | 47 | 62 | 65 | 65 | 3498 | 0.006 | 2 | 38 | 62 | 65 | Y | Y |
| R | 85 | 161 | 10627 | 0.02 | 3 | 18 | 109 | 160 | 85 | 61 | 3820 | 0.006 | 1 | 32 | 56 | 61 | Y | N |
| S | 205 | 529 | 49330 | 0.08 | 4 | 56 | 234 | 495 | 205 | 293 | 22767 | 0.04 | 9 | 122 | 255 | 290 | Y | Y |
| T | 279 | 290 | 29359 | 0.05 | 1 | 25 | 126 | 272 | 278 | 157 | 13902 | 0.02 | 3 | 58 | 138 | 154 | N | N |
| U | 354 | 614 | 56972 | 0.09 | 9 | 70 | 293 | 585 | 354 | 306 | 25381 | 0.04 | 6 | 147 | 269 | 306 | N | N |
| V | 161 | 961 | 83655 | 0.13 | 7 | 87 | 489 | 943 | 161 | 461 | 35217 | 0.06 | 16 | 204 | 403 | 461 | Y | N |

<sup>a</sup> Analysis performed with only with 21 mtDNA blocks. Cytochromes gene sequences were not included in the repeat library.
<sup>b</sup> Insertion may not be of the entire element.

**Table S3.**
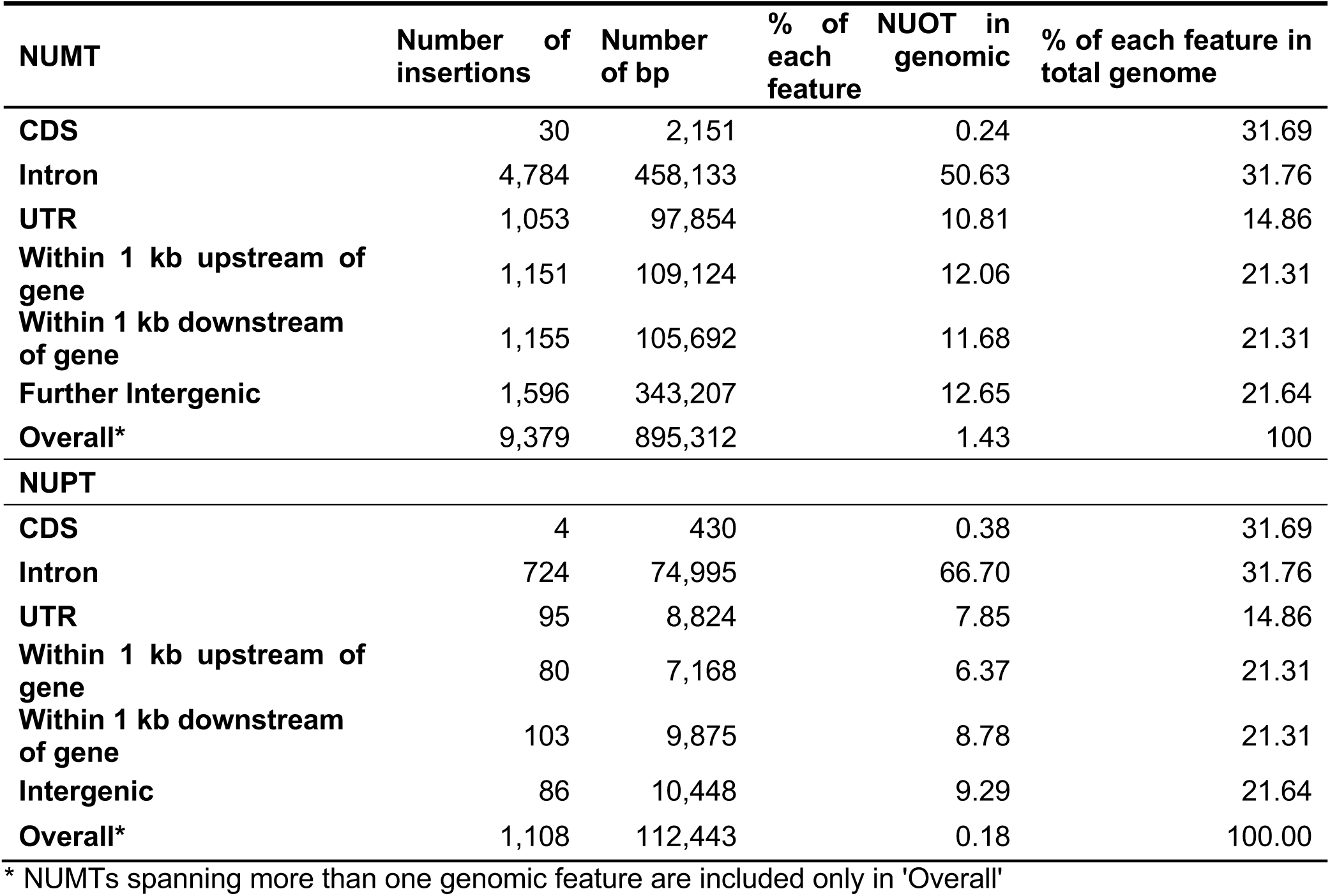
Categorization of NUOTs by *T. gondii* ME49 genome feature

**Table S4.**
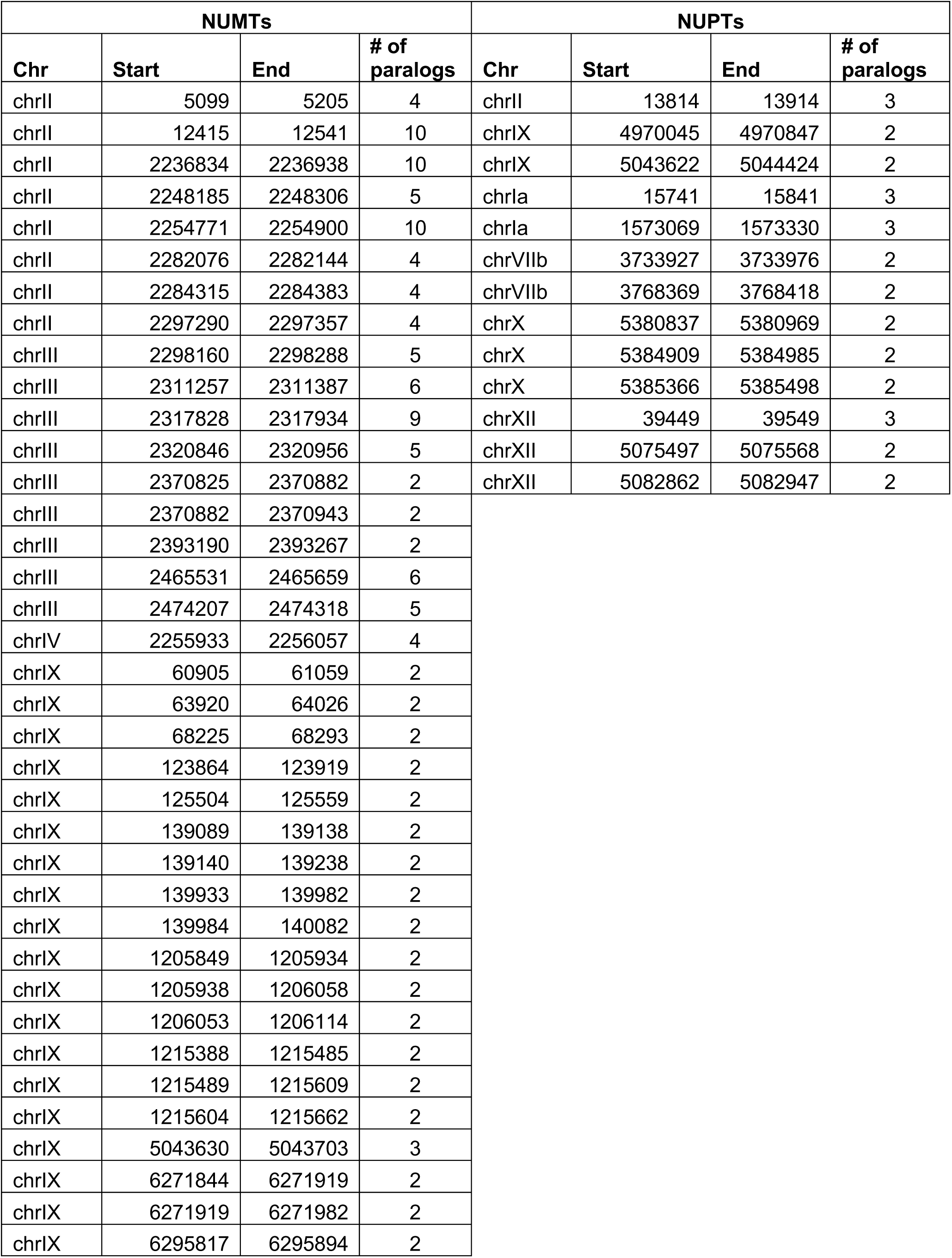
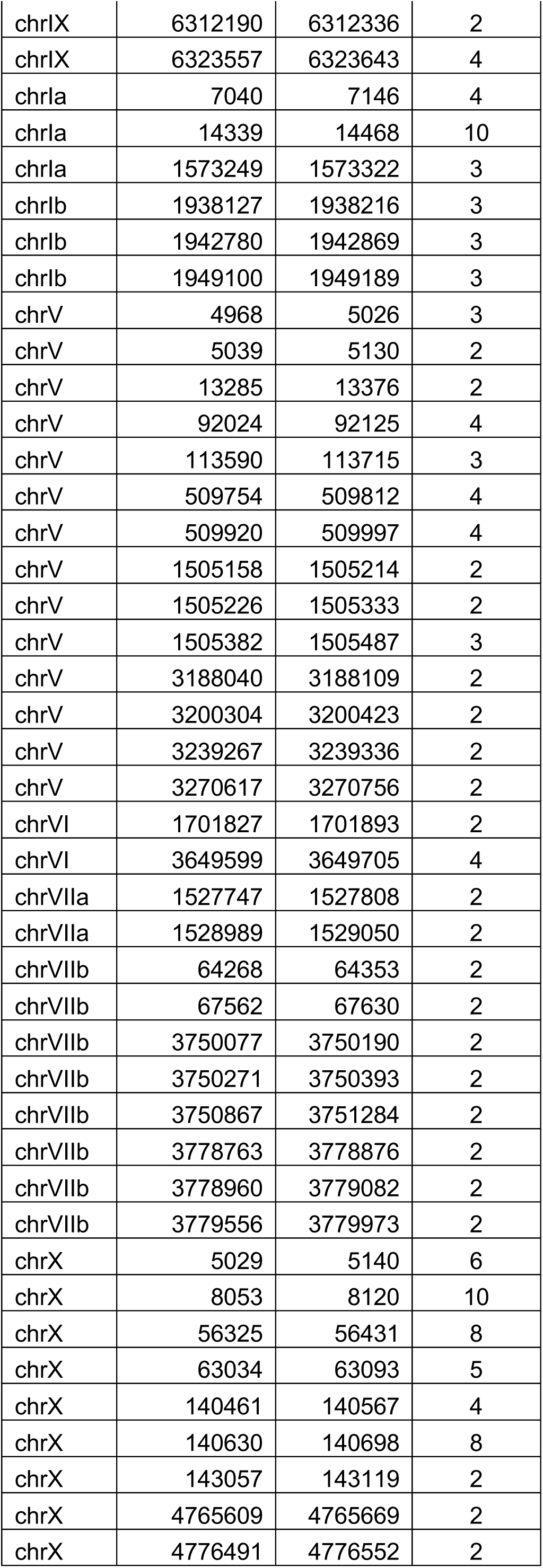
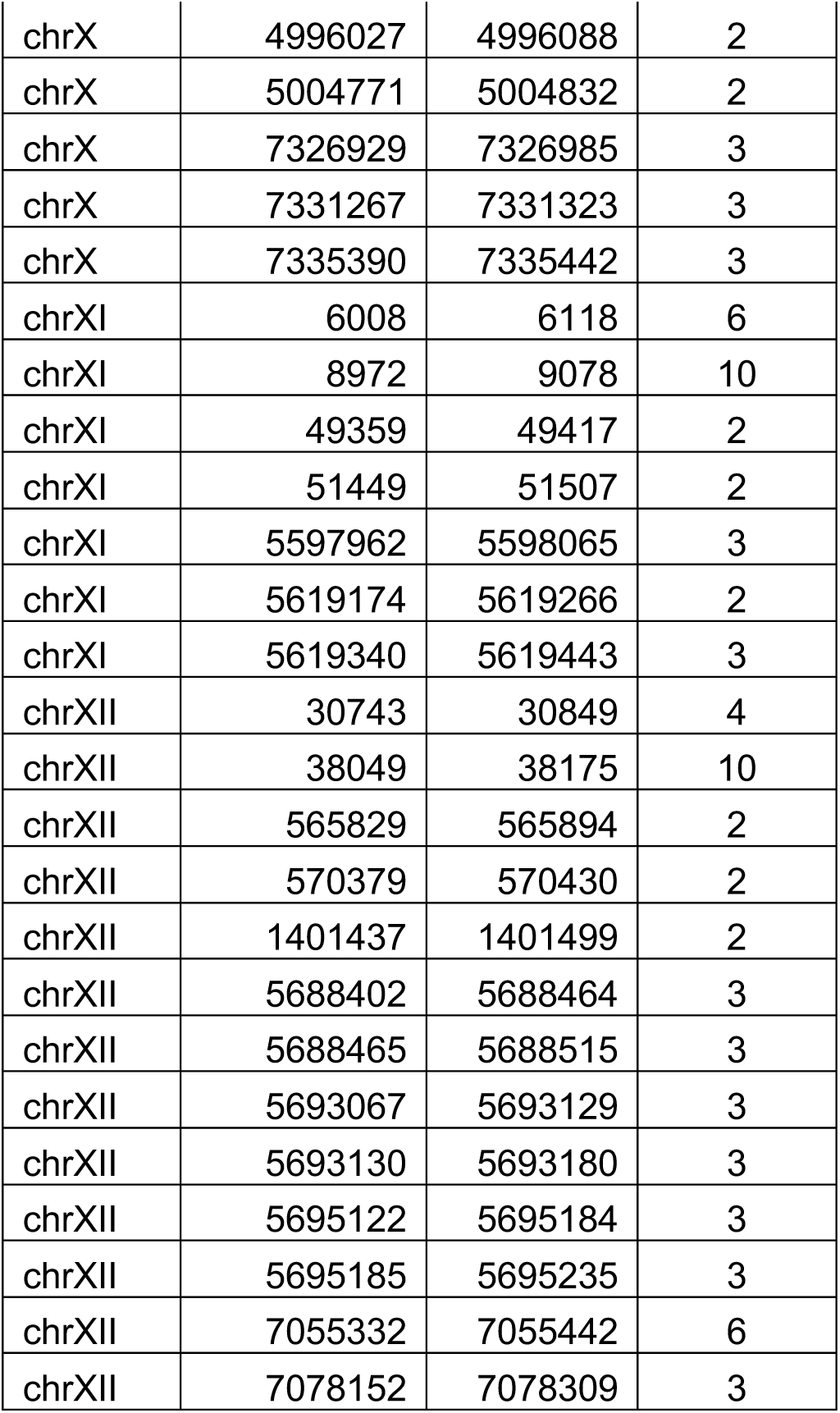
NUM/PTs that show duplication in the nuclear genome sequence

| NUMTs |  |  |  | NUPTs |  |  |  |
| --- | --- | --- | --- | --- | --- | --- | --- |
| Chr | Start | End | # of paralogs | Chr | Start | End | # of paralogs |
| chrII | 5099 | 5205 | 4 | chrII | 13814 | 13914 | 3 |
| chrII | 12415 | 12541 | 10 | chrIX | 4970045 | 4970847 | 2 |
| chrII | 2236834 | 2236938 | 10 | chrIX | 5043622 | 5044424 | 2 |
| chrII | 2248185 | 2248306 | 5 | chrIa | 15741 | 15841 | 3 |
| chrII | 2254771 | 2254900 | 10 | chrIa | 1573069 | 1573330 | 3 |
| chrII | 2282076 | 2282144 | 4 | chrVIIb | 3733927 | 3733976 | 2 |
| chrII | 2284315 | 2284383 | 4 | chrVIIb | 3768369 | 3768418 | 2 |
| chrII | 2297290 | 2297357 | 4 | chrX | 5380837 | 5380969 | 2 |
| chrIII | 2298160 | 2298288 | 5 | chrX | 5384909 | 5384985 | 2 |
| chrIII | 2311257 | 2311387 | 6 | chrX | 5385366 | 5385498 | 2 |
| chrIII | 2317828 | 2317934 | 9 | chrXII | 39449 | 39549 | 3 |
| chrIII | 2320846 | 2320956 | 5 | chrXII | 5075497 | 5075568 | 2 |
| chrIII | 2370825 | 2370882 | 2 | chrXII | 5082862 | 5082947 | 2 |
| chrIII | 2370882 | 2370943 | 2 |  |  |  |  |
| chrIII | 2393190 | 2393267 | 2 |  |  |  |  |
| chrIII | 2465531 | 2465659 | 6 |  |  |  |  |
| chrIII | 2474207 | 2474318 | 5 |  |  |  |  |
| chrIV | 2255933 | 2256057 | 4 |  |  |  |  |
| chrIX | 60905 | 61059 | 2 |  |  |  |  |
| chrIX | 63920 | 64026 | 2 |  |  |  |  |
| chrIX | 68225 | 68293 | 2 |  |  |  |  |
| chrIX | 123864 | 123919 | 2 |  |  |  |  |
| chrIX | 125504 | 125559 | 2 |  |  |  |  |
| chrIX | 139089 | 139138 | 2 |  |  |  |  |
| chrIX | 139140 | 139238 | 2 |  |  |  |  |
| chrIX | 139933 | 139982 | 2 |  |  |  |  |
| chrIX | 139984 | 140082 | 2 |  |  |  |  |
| chrIX | 1205849 | 1205934 | 2 |  |  |  |  |
| chrIX | 1205938 | 1206058 | 2 |  |  |  |  |
| chrIX | 1206053 | 1206114 | 2 |  |  |  |  |
| chrIX | 1215388 | 1215485 | 2 |  |  |  |  |
| chrIX | 1215489 | 1215609 | 2 |  |  |  |  |
| chrIX | 1215604 | 1215662 | 2 |  |  |  |  |
| chrIX | 5043630 | 5043703 | 3 |  |  |  |  |
| chrIX | 6271844 | 6271919 | 2 |  |  |  |  |
| chrIX | 6271919 | 6271982 | 2 |  |  |  |  |
| chrIX | 6295817 | 6295894 | 2 |  |  |  |  |

|  |  |  |  |
| --- | --- | --- | --- |
| chrIX | 6312190 | 6312336 | 2 |
| chrIX | 6323557 | 6323643 | 4 |
| chrIa | 7040 | 7146 | 4 |
| chrIa | 14339 | 14468 | 10 |
| chrIa | 1573249 | 1573322 | 3 |
| chrIb | 1938127 | 1938216 | 3 |
| chrIb | 1942780 | 1942869 | 3 |
| chrIb | 1949100 | 1949189 | 3 |
| chrV | 4968 | 5026 | 3 |
| chrV | 5039 | 5130 | 2 |
| chrV | 13285 | 13376 | 2 |
| chrV | 92024 | 92125 | 4 |
| chrV | 113590 | 113715 | 3 |
| chrV | 509754 | 509812 | 4 |
| chrV | 509920 | 509997 | 4 |
| chrV | 1505158 | 1505214 | 2 |
| chrV | 1505226 | 1505333 | 2 |
| chrV | 1505382 | 1505487 | 3 |
| chrV | 3188040 | 3188109 | 2 |
| chrV | 3200304 | 3200423 | 2 |
| chrV | 3239267 | 3239336 | 2 |
| chrV | 3270617 | 3270756 | 2 |
| chrVI | 1701827 | 1701893 | 2 |
| chrVI | 3649599 | 3649705 | 4 |
| chrVIIa | 1527747 | 1527808 | 2 |
| chrVIIa | 1528989 | 1529050 | 2 |
| chrVIIb | 64268 | 64353 | 2 |
| chrVIIb | 67562 | 67630 | 2 |
| chrVIIb | 3750077 | 3750190 | 2 |
| chrVIIb | 3750271 | 3750393 | 2 |
| chrVIIb | 3750867 | 3751284 | 2 |
| chrVIIb | 3778763 | 3778876 | 2 |
| chrVIIb | 3778960 | 3779082 | 2 |
| chrVIIb | 3779556 | 3779973 | 2 |
| chrX | 5029 | 5140 | 6 |
| chrX | 8053 | 8120 | 10 |
| chrX | 56325 | 56431 | 8 |
| chrX | 63034 | 63093 | 5 |
| chrX | 140461 | 140567 | 4 |
| chrX | 140630 | 140698 | 8 |
| chrX | 143057 | 143119 | 2 |
| chrX | 4765609 | 4765669 | 2 |
| chrX | 4776491 | 4776552 | 2 |
| chrX | 4996027 | 4996088 | 2 |
| chrX | 5004771 | 5004832 | 2 |
| chrX | 7326929 | 7326985 | 3 |
| chrX | 7331267 | 7331323 | 3 |
| chrX | 7335390 | 7335442 | 3 |
| chrXI | 6008 | 6118 | 6 |
| chrXI | 8972 | 9078 | 10 |
| chrXI | 49359 | 49417 | 2 |
| chrXI | 51449 | 51507 | 2 |
| chrXI | 5597962 | 5598065 | 3 |
| chrXI | 5619174 | 5619266 | 2 |
| chrXI | 5619340 | 5619443 | 3 |
| chrXII | 30743 | 30849 | 4 |
| chrXII | 38049 | 38175 | 10 |
| chrXII | 565829 | 565894 | 2 |
| chrXII | 570379 | 570430 | 2 |
| chrXII | 1401437 | 1401499 | 2 |
| chrXII | 5688402 | 5688464 | 3 |
| chrXII | 5688465 | 5688515 | 3 |
| chrXII | 5693067 | 5693129 | 3 |
| chrXII | 5693130 | 5693180 | 3 |
| chrXII | 5695122 | 5695184 | 3 |
| chrXII | 5695185 | 5695235 | 3 |
| chrXII | 7055332 | 7055442 | 6 |
| chrXII | 7078152 | 7078309 | 3 |

**Table S5.** Strain-specific NUMTs in ME49 GT1 and VEG

| **Coordinates of NUMTs<sup>a</sup>** | **Divergence (%)** | **Associated nuclear gene** | **Location in gene** | **Only present /absent in** | **Insertion/deletion** |
| --- | --- | --- | --- | --- | --- |
| TGGT1_chrII:<br>244462..244554 | 4.4 | TGGT1_221330 | intron | GT1 | insertion |
| TGGT1_chrII:<br>1102510..1102622 | 27.4 | TGGT1_222410 | 1.8 Kb upstream | ME49 | deletion |
| TGME49_chrIII:<br>1730617..1730859 | 23.5 | TGME49_254420 | intron | ME49 | not sure |
| TGME49_chrIV:<br>1102974..1103065 | 20.6 | TGME49_318770 | 1.3 Kb upstream | VEG | deletion |
| TGME49_chrIX:<br>2923871..2923932 | 3.2 | none | none | ME49 | insertion |
| TGME49_chrIX:<br>3047280..3047543 | 12.5 | TGME49_289180 | intron | VEG | deletion |
| TGGT1_chrVI:<br>699247..699347 | 1 | TGGT1_239480 | intron | GT1 | insertion |
| TGGT1_chrVIII:<br>3463130..3463400 | 24.5 | TGGT1_273478 | intron | GT1 | not sure |
| TGME49_chrVIII:<br>4435556..4435633 | 20.8 | none | none | ME49 | not sure |
| TGGT1_chrVIIa:<br>2687959..2688077 | 7.6 | TGGT1_203135 | intron | VEG | not sure |
| TGME49_chrVIIa:<br>3250536..3250697 | 30.6 | TGME49_202540 | 1.3 Kb upstream | VEG | deletion |
| TGGT1_chrVIIb:<br>1000247..1000321 | 14.7 | TGGT1_262780 | 173 bp downstream | ME49 | deletion |
| TGGT1_chrVIIb:<br>3183923..3184103 | 23.2 | none | none | GT1 | not sure |
| TGGT1_chrVIIb:<br>4560651..4560737 | 21.2 | TGGT1_255890 | intron | GT1 | not sure |
| TGME49_chrX:<br>475518..475568 | 23.5 | TGME49_228230 | 3' UTR | ME49 | not sure |
| TGGT1_chrX:<br>2610785..2610846 | 17.7 | TGGT1_224900 | 591 bp upstream | ME49 | deletion |
| TGGT1_chrX:<br>2885685..2885842 | 24.1 | TGGT1_224540B | intron | ME49 | deletion |
| TGME49_chrX:<br>3011523..3011876 | 16.9 | TGME49_224510 | 257 bp downstream | VEG | deletion |
| TGME49_chrX:<br>6399715..6399810 | 21.3 | TGME49_014600 | intron | VEG | deletion |
| TGGT1_chrX:<br>6755042..6755100 | 25.4 | TGGT1_215260 | intron | GT1 | not sure |
| TGGT1_chrXI:<br>4963753..4963855 | 25.2 | none | none | GT1 | not sure |
| TGGT1_chrXII:<br>287472..287613 | 22.7 | TGGT1_299820 | 1.1 Kb upstream | ME49 | not sure |
| TGGT1_chrXII:<br>2346630..2346730 | 21.7 | none | none | GT1 | not sure |
| TGGT1_chrXII:<br>2367574..2367660 | 25.3 | TGGT1_245610 | intron | GT1 | not sure |
| TGME49_chrVIII:<br>2496663..2496886 | 19.1 | TGME49_233200 | 1.1 Kb<br>upstream | GT1 | deletion |
| TGME49_chrVIIa:<br>3199077..3199253 | 22.5 | TGME49_202580 | intron | VEG | deletion |
| TGME49_chrVIIb:<br>701168..701236 | 24.6 | TGME49_263220 | intron | ME49 | not sure |
| TGME49_chrVIIb:<br>1710367..1710510 | 25 | TGME49_261490 | intron | ME49 | not sure |
| TGME49_chrXI:<br>1310258..1310322 | 24.6 | TGME49_310380 | intron | GT1 | deletion |
| TGME49_chrXI:<br>3550511..3550993 | 19.4 | TGME49_313620 | 400 bp<br>upstream | GT1 | deletion |
| TGME49_chrXII:<br>3864844..3864916 | 0 | TGME49_248450 | intron | GT1 | not sure |

<sup>a</sup>Includes NUMTs that are partially present or absent in the other strain

**Table S6.**
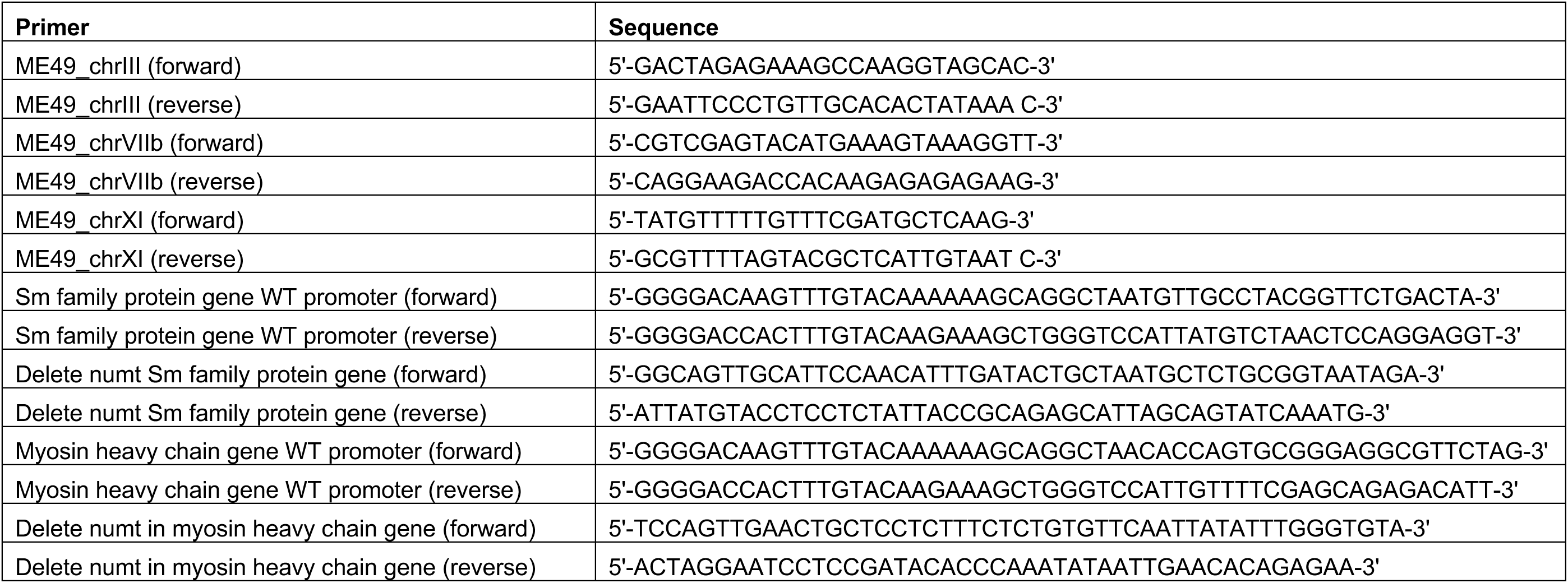
Primers used in Figure S1 and to amplify and alter the promoters used in the transfection assays (Fig. 3)

**Table S7.** NUMT content in *T. gondii* strains

| **Species** | **Density (%)** | **Number of insertions** | **Number of base pairs** |
| --- | --- | --- | --- |
| *T. gondii* GT1 | 1.44 | 9 175 | 874 525 |
| *T. gondii* VEG | 1.41 | 9 270 | 883 437 |
| *T. gondii* CAST | 1.39 | 9 370 | 876 895 |
| *T. gondii* TgCtCo5 | 1.4 | 9 411 | 874 358 |
| *T. gondii* TgCATBr9 | 1.42 | 9 332 | 874 826 |
| *T. gondii* GAB2 | 1.4 | 9 373 | 880 313 |
| *T. gondii* FOU | 1.41 | 9 314 | 871 351 |
| *T. gondii* TgCATBr5 | 1.41 | 9 370 | 869 048 |
| *T. gondii* MAS | 1.42 | 9 318 | 874 616 |
| *T. gondii* p89 | 1.41 | 9 314 | 875 326 |
| *T. gondii* ME49 | 1.43 | 9 379 | 895 312 |
| *T. gondii* ARI | 1.4 | 9 445 | 883 577 |
| *T. gondii* TgCat_PRC2 | 1.4 | 9 485 | 880 946 |
| *T. gondii* COUG | 1.39 | 9 525 | 884 389 |
| *T. gondii* VAND | 1.41 | 9 331 | 879 564 |
| *T. gondii* RUB | 1.41 | 9 426 | 880 485 |

**Table S8.** Location of NUMTs displaying differential presence/absence in 16 *T. gondii* strains

| Numt #<sup>s</sup> | Length (bp) | % Div | ME49 chr | Start* | End* | CAST | TgCTCo5 | GT1 | TgCATBr9 | GAB2 | FOU | TgCATBr5 | MAS | p89 | VEG | ME49 | ARI | TgCat_PRC2 | COUG | VAND | RUB | Location* | Associated gene |
| --- | --- | --- | --- | --- | --- | --- | --- | --- | --- | --- | --- | --- | --- | --- | --- | --- | --- | --- | --- | --- | --- | --- | --- |
| 1 | 61 | 14 | chrIa | 1018474 | 1018944 | 1 | 1 | 1 | 1 | 1 | 1 | 1 | 1 | 1 | 1 | 1 | 1 | 1 | 1 | 0 | 0 | I | 294630 |
| 2 | 73 | 20 | chrIb | 213457 | 213857 | 0 | 0 | 0 | 0 | 0 | 0 | 0 | 1 | 1 | 0 | 0 | 1 | 1 | 1 | 0 | 1 | UF | 207690 |
| 3 | 67 | 7.7 | chrIb | 1596582 | 1596989 | 1 | 0 | 1 | 0 | 0 | 0 | 0 | 1 | 0 | 0 | 0 | 0 | 0 | 1 | 0 | 0 | I | 209970 |
| 4 | 45 | 2.3 | chrII | 261484 | 261902 | 0 | 0 | 1 | 1 | 0 | 1 | 1 | 0 | 0 | 0 | 0 | 0 | 0 | 0 | 1 | 1 | I | 221330 |
| 5 | 67 | 1.5 | chrIII | 2457884 | 2458465 | 1 | 1 | 1 | 0 | 1 | 1 | 1 | 1 | 1 | 1 | 1 | 1 | 1 | 1 | 1 | 1 | IG |  |
| 6 | 28 | 0 | chrIV | 1660519 | 1660918 | 0 | 0 | 0 | 0 | 0 | 0 | 0 | 1 | 0 | 0 | 0 | 0 | 0 | 0 | 0 | 0 | E | 211630 |
| 7 | 157 | 1.9 | chrIV | 2006407 | 2006806 | 0 | 0 | 0 | 1 | 0 | 0 | 1 | 1 | 0 | 0 | 0 | 0 | 0 | 0 | 0 | 0 | DF | 210960 |
| 8 | 63 | 6.2 | chrV | 1768829 | 1769234 | 0 | 0 | 0 | 0 | 0 | 0 | 0 | 0 | 0 | 0 | 0 | 0 | 1 | 0 | 0 | 0 | E | 286590 |
| 9 | 51 | 16 | chrV | 2529301 | 2529857 | 0 | 0 | 0 | 0 | 0 | 0 | 0 | 0 | 0 | 0 | 1 | 1 | 0 | 1 | 0 | 0 | I | 285210 |
| 10 | 52 | 0 | chrV | 2798255 | 2798857 | 1 | 1 | 1 | 1 | 1 | 1 | 0 | 0 | 1 | 1 | 0 | 0 | 0 | 0 | 1 | 1 | I | 284170 |
| 11 | 100 | 1 | chrVI | 706677 | 707077 | 1 | 1 | 1 | 1 | 1 | 1 | 1 | 0 | 1 | 0 | 0 | 0 | 0 | 0 | 1 | 1 | I | 239480 |
| 12 | 91 | 23.3 | chrVI | 2259261 | 2259751 | 1 | 1 | 1 | 1 | 1 | 1 | 1 | 0 | 1 | 1 | 1 | 1 | 1 | 1 | 1 | 1 | E | 242770 |
| 13 | 64 | 0 | chrVI | 2415880 | 2416280 | 0 | 0 | 0 | 0 | 0 | 0 | 0 | 0 | 1 | 0 | 0 | 0 | 0 | 0 | 1 | 0 | UF | 243140 |
| 14 | 88 | 26.1 | chrVIIa | 605108 | 605597 | 1 | 1 | 1 | 0 | 1 | 1 | 0 | 1 | 1 | 1 | 1 | 1 | 1 | 1 | 1 | 1 | I | 304650 |
| 15 | 133 | 27.8 | chrVIIa | 2259268 | 2259788 | 1 | 0 | 1 | 1 | 1 | 1 | 1 | 1 | 0 | 0 | 1 | 1 | 1 | 1 | 1 | 1 | I | 203760 |
| 16 | 54 | 14.6 | chrVIIa | 2673706 | 2674160 | 1 | 1 | 0 | 1 | 0 | 0 | 1 | 1 | 1 | 1 | 1 | 1 | 1 | 1 | 1 | 0 | DF | 203220 |
| 17 | 162 | 30.6 | chrVIIa | 3250336 | 3250897 | 1 | 1 | 1 | 1 | 1 | 1 | 1 | 1 | 1 | 0 | 1 | 1 | 1 | 1 | 0 | 1 | E | 202540 |
| 18 | 59 | 0 | chrVIIb | 668720 | 669119 | 0 | 0 | 0 | 0 | 0 | 0 | 1 | 1 | 0 | 0 | 0 | 0 | 0 | 0 | 0 | 0 | I | 263270 |
| 19 | 75 | 14.7 | chrVIIb | 1025866 | 1026297 | 1 | 1 | 1 | 1 | 1 | 1 | 1 | 1 | 1 | 1 | 0 | 0 | 1 | 0 | 1 | 1 | I | 262780 |
| 20 | 140 | 26.5 | chrVIIb | 1710159 | 1710710 | 0 | 0 | 0 | 0 | 0 | 0 | 0 | 0 | 0 | 0 | 1 | 1 | 1 | 1 | 0 | 0 | I | 261490 |
| 21 | 53 | 1.9 | chrVIIb | 1904265 | 1904665 | 0 | 0 | 0 | 0 | 0 | 0 | 0 | 0 | 0 | 0 | 0 | 0 | 1 | 1 | 0 | 0 | IG |  |
| 22 | 181 | 23.2 | chrVIIb | 3218491 | 3219027 | 1 | 1 | 1 | 1 | 1 | 1 | 1 | 1 | 1 | 0 | 0 | 0 | 1 | 1 | 1 | 1 | IG |  |
| 23 | 87 | 21.2 | chrVIIb | 4646913 | 4647351 | 1 | 1 | 1 | 1 | 1 | 1 | 1 | 1 | 1 | 0 | 0 | 0 | 1 | 0 | 1 | 1 | I | 255890 |
| 24 | 77 | 22.1 | chrVIII | 2496611 | 2497086 | 1 | 0 | 0 | 0 | 0 | 0 | 0 | 1 | 1 | 1 | 1 | 1 | 1 | 1 | 0 | 1 | IG |  |
| 25 | 99 | 23.2 | chrVIII | 3466455 | 3466854 | 0 | 1 | 1 | 1 | 1 | 1 | 0 | 1 | 0 | 0 | 0 | 0 | 0 | 0 | 0 | 1 | I | 243482 |
| 26 | 56 | 14.3 | chrVIII | 4435378 | 4435833 | 0 | 0 | 0 | 0 | 0 | 0 | 0 | 1 | 0 | 0 | 1 | 1 | 1 | 1 | 0 | 0 | DF | 271740 |
| 27 | 100 | 28.6 | chrIX | 3008823 | 3009336 | 1 | 1 | 1 | 1 | 1 | 1 | 1 | 1 | 1 | 1 | 1 | 1 | 1 | 0 | 1 | 1 | I | 289120 |
| 28 | 133 | 29.3 | chrIX | 3038491 | 3039017 | 1 | 1 | 1 | 0 | 1 | 1 | 0 | 0 | 1 | 1 | 1 | 1 | 1 | 1 | 1 | 1 | I | 289170 |
| 29 | 73 | 20.6 | chrIX | 4251589 | 4252066 | 1 | 1 | 1 | 0 | 1 | 1 | 0 | 0 | 1 | 1 | 1 | 1 | 1 | 1 | 1 | 1 | IG |  |
| 30 | 51 | 23.5 | chrX | 475318 | 475768 | 0 | 0 | 0 | 0 | 0 | 0 | 0 | 1 | 0 | 0 | 1 | 1 | 1 | 1 | 0 | 0 | E | 228230 |
| 31 | 162 | 17.7 | chrX | 2699533 | 2699941 | 1 | 1 | 1 | 1 | 0 | 1 | 1 | 1 | 1 | 1 | 0 | 0 | 1 | 1 | 1 | 1 | UF | 224900 |
| 32 | 81 | 21.9 | chrX | 2977559 | 2977959 | 1 | 1 | 1 | 1 | 0 | 1 | 1 | 1 | 1 | 1 | 0 | 1 | 1 | 1 | 1 | 1 | I | 224540 |
| 33 | 74 | 17.6 | chrX | 3011324 | 3011796 | 1 | 1 | 1 | 1 | 1 | 1 | 0 | 1 | 0 | 0 | 1 | 0 | 1 | 1 | 0 | 1 | I | 224510 |
| 34 | 52 | 13.5 | chrX | 5158557 | 5159008 | 1 | 1 | 1 | 1 | 1 | 1 | 1 | 1 | 1 | 1 | 1 | 1 | 1 | 1 | 0 | 1 | I | 235920 |
| 35 | 78 | 0 | chrX | 6191403 | 6191803 | 0 | 1 | 0 | 0 | 0 | 0 | 0 | 0 | 0 | 1 | 0 | 0 | 0 | 0 | 1 | 0 | I | 214260 |
| 36 | 84 | 23.2 | chrX | 6399526 | 6400012 | 1 | 1 | 1 | 1 | 1 | 1 | 1 | 0 | 0 | 0 | 1 | 1 | 1 | 1 | 1 | 1 | I | 214600 |
| 37 | 59 | 0 | chrX | 6727704 | 6728163 | 0 | 0 | 0 | 0 | 1 | 0 | 0 | 0 | 0 | 0 | 1 | 0 | 0 | 0 | 0 | 0 | UF | 215060 |
| 38 | 58 | 25.4 | chrX | 6863803 | 6864202 | 0 | 0 | 1 | 0 | 0 | 0 | 1 | 0 | 1 | 0 | 0 | 0 | 0 | 0 | 1 | 0 | I | 215260 |
| 39 | 166 | 19.4 | chrXI | 1532798 | 1533403 | 1 | 1 | 1 | 1 | 1 | 1 | 1 | 1 | 0 | 1 | 1 | 1 | 1 | 1 | 1 | 1 | DF | 310710 |
| 40 | 239 | 25.8 | chrXI | 1603990 | 1604629 | 1 | 1 | 1 | 0 | 0 | 0 | 1 | 1 | 1 | 1 | 1 | 1 | 1 | 1 | 1 | 1 | DF | 310802 |
| 41 | 85 | 2.4 | chrXI | 2153288 | 2153772 | 0 | 0 | 1 | 1 | 0 | 1 | 1 | 1 | 1 | 0 | 1 | 0 | 0 | 0 | 1 | 1 | IG |  |
| 42 | 48 | 0 | chrXI | 2168335 | 2168735 | 0 | 0 | 0 | 0 | 0 | 0 | 1 | 1 | 0 | 0 | 0 | 0 | 0 | 0 | 0 | 0 | I | 311625 |
| 43 | 57 | 5.2 | chrXI | 2292428 | 2293114 | 0 | 0 | 0 | 0 | 0 | 0 | 1 | 1 | 0 | 0 | 0 | 0 | 0 | 0 | 0 | 0 | E | 311800 |
| 44 | 102 | 24 | chrXI | 2928696 | 2929196 | 0 | 1 | 1 | 0 | 0 | 0 | 1 | 1 | 1 | 1 | 1 | 1 | 1 | 1 | 1 | 1 | I | 312820 |
| 45 | 37 | 0 | chrXI | 3138783 | 3139183 | 0 | 0 | 0 | 1 | 1 | 1 | 0 | 0 | 0 | 0 | 0 | 0 | 0 | 0 | 0 | 0 | E | 313150 |
| 46 | 122 | 24.1 | chrXI | 3380178 | 3380628 | 1 | 0 | 0 | 1 | 1 | 1 | 1 | 1 | 1 | 0 | 0 | 0 | 1 | 1 | 1 | 1 | I | 313430 |
| 47 | 58 | 12.1 | chrXI | 3550551 | 3551008 | 0 | 0 | 0 | 0 | 0 | 0 | 0 | 0 | 0 | 1 | 1 | 0 | 0 | 0 | 0 | 0 | I | 313630 |
| 48 | 103 | 25.2 | chrXI | 5020102 | 5020569 | 1 | 0 | 1 | 1 | 0 | 1 | 1 | 1 | 1 | 0 | 0 | 0 | 0 | 0 | 0 | 0 | I | 315845 |
| 49 | 38 | 2.6 | chrXI | 5133457 | 5133946 | 0 | 1 | 0 | 1 | 1 | 1 | 0 | 1 | 0 | 1 | 1 | 1 | 1 | 1 | 0 | 0 | E | 316100 |
| 50 | 142 | 22.7 | chrXII | 328834 | 329234 | 1 | 1 | 1 | 1 | 1 | 1 | 1 | 1 | 1 | 1 | 0 | 1 | 1 | 1 | 1 | 1 | UF | 299820 |
| 51 | 84 | 1.2 | chrXII | 905189 | 905588 | 0 | 0 | 0 | 1 | 1 | 1 | 1 | 0 | 1 | 1 | 0 | 0 | 0 | 0 | 0 | 0 | I | 219500 |
| 52 | 79 | 25 | chrXII | 2430105 | 2430505 | 1 | 1 | 1 | 1 | 0 | 1 | 1 | 1 | 1 | 0 | 0 | 0 | 0 | 1 | 1 | 1 | I | 245610 |
| 53 | 73 | 0 | chrXII | 3864644 | 3865116 | 0 | 0 | 0 | 0 | 0 | 0 | 0 | 0 | 0 | 1 | 1 | 1 | 0 | 1 | 0 | 0 | I | 248450 |
| 54 | 60 | 20 | chrXII | 4009932 | 4010391 | 0 | 0 | 0 | 0 | 1 | 1 | 0 | 0 | 1 | 1 | 1 | 1 | 1 | 1 | 1 | 0 | UF | 248580 |
| 55 | 35 | 0 | chrXII | 4599466 | 4599866 | 0 | 1 | 0 | 0 | 1 | 1 | 0 | 1 | 1 | 0 | 0 | 0 | 1 | 0 | 1 | 0 | I | 249530 |
| 56 | 41 | 13.7 | chrXII | 4641543 | 4642129 | 1 | 0 | 1 | 1 | 0 | 0 | 1 | 1 | 0 | 1 | 1 | 1 | 1 | 1 | 1 | 1 | I | 249560 |

<sup>§</sup>Only NUMTs that are present/absent almost entirely are included.
\* Location of NUMT is with respect to the ME49 genome sequence. Start and stop co-ordinates include 200 bp up and downstream of NUMT. Genomic locations are indicated as Exon = E Intron = I Upstream flank = UF downstream flank = DF Intergenic = IG (intergenic does not include flanking regions). Coordinates for NUMT location include 200 bp upstream and downstream flanking regions of the NUMT. Presence =1 absence 0.

**Table S9.** Enrichment analyses of genes associated with NUMTs displaying differential presence/absence

| **ID** | **Name** | **Bgd count** | **Result count** | **Result gene list** | **Pct of bgd** | **Fold enrichment** | **Odds ratio** | **P-value** | **Benjamini** | **Bonferroni** |
| --- | --- | --- | --- | --- | --- | --- | --- | --- | --- | --- |
| **GO:0007017** | microtubule-based process | 72 | 4 | TGME49_209970<br>TGME49_235920<br>TGME49_243482<br>TGME49_315845 | 5.6 | 7.95 | 9.34 | 0.001465 | 0.18598 | 0.221162 |
| **GO:0007018** | microtubule-based movement | 46 | 3 | TGME49_235920<br>TGME49_243482<br>TGME49_315845 | 6.5 | 9.33 | 10.77 | 0.003861 | 0.18598 | 0.583045 |
| **GO:0006928** | movement of cell or subcellular component | 49 | 3 | TGME49_235920<br>TGME49_243482<br>TGME49_315845 | 6.1 | 8.76 | 10.06 | 0.00462 | 0.18598 | 0.69761 |
| **GO:0006071** | glycerol metabolic process | 2 | 1 | TGME49_263270 | 50 | 71.51 | 146.3 | 0.013936 | 0.18598 | 1 |
| **GO:0046116** | queuosine metabolic process | 2 | 1 | TGME49_316100 | 50 | 71.51 | 146.3 | 0.013936 | 0.18598 | 1 |
| **GO:0019400** | alditol metabolic process | 2 | 1 | TGME49_263270 | 50 | 71.51 | 146.3 | 0.013936 | 0.18598 | 1 |
| **GO:0008616** | queuosine biosynthetic process | 2 | 1 | TGME49_316100 | 50 | 71.51 | 146.3 | 0.013936 | 0.18598 | 1 |
| **GO:0023052** | signaling | 84 | 3 | TGME49_202540<br>TGME49_215060<br>TGME49_289170 | 3.6 | 5.11 | 5.67 | 0.020239 | 0.18598 | 1 |
| **GO:0007165** | signal transduction | 84 | 3 | TGME49_202540<br>TGME49_215060<br>TGME49_289170 | 3.6 | 5.11 | 5.67 | 0.020239 | 0.18598 | 1 |
| **GO:0055086** | nucleobase-containing small molecule metabolic process | 85 | 3 | TGME49_215260<br>TGME49_289170<br>TGME49_316100 | 3.5 | 5.05 | 5.6 | 0.02088 | 0.18598 | 1 |
| **GO:0018130** | heterocycle biosynthetic process | 153 | 4 | TGME49_215260<br>TGME49_249560<br>TGME49_289170<br>TGME49_316100 | 2.6 | 3.74 | 4.19 | 0.020974 | 0.18598 | 1 |
| **GO:0007154** | cell communication | 86 | 3 | TGME49_202540<br>TGME49_215060<br>TGME49_289170 | 3.5 | 4.99 | 5.53 | 0.021533 | 0.18598 | 1 |
| **GO:1901362** | organic cyclic compound biosynthetic process | 158 | 4 | TGME49_215260<br>TGME49_249560<br>TGME49_289170<br>TGME49_316100 | 2.5 | 3.62 | 4.05 | 0.023304 | 0.18598 | 1 |
| **GO:0006207** | 'de novo' pyrimidine nucleobase biosynthetic process | 4 | 1 | TGME49_215260 | 25 | 35.76 | 48.75 | 0.027683 | 0.18598 | 1 |
| **GO:0051716** | cellular response to stimulus | 170 | 4 | TGME49_202540<br>TGME49_210960<br>TGME49_215060<br>TGME49_289170 | 2.4 | 3.37 | 3.75 | 0.029529 | 0.18598 | 1 |
| **GO:0046785** | microtubule polymerization | 5 | 1 | TGME49_209970 | 20 | 28.61 | 36.55 | 0.034486 | 0.18598 | 1 |
| **GO:0046112** | nucleobase biosynthetic process | 5 | 1 | TGME49_215260 | 20 | 28.61 | 36.55 | 0.034486 | 0.18598 | 1 |
| **GO:0042455** | ribonucleoside biosynthetic process | 5 | 1 | TGME49_316100 | 20 | 28.61 | 36.55 | 0.034486 | 0.18598 | 1 |
| **GO:0009112** | nucleobase metabolic process | 5 | 1 | TGME49_215260 | 20 | 28.61 | 36.55 | 0.034486 | 0.18598 | 1 |
| **GO:0034404** | nucleobase-containing small molecule biosynthetic process | 5 | 1 | TGME49_316100 | 20 | 28.61 | 36.55 | 0.034486 | 0.18598 | 1 |
| **GO:1901659** | glycosyl compound biosynthetic process | 5 | 1 | TGME49_316100 | 20 | 28.61 | 36.55 | 0.034486 | 0.18598 | 1 |
| **GO:0007020** | microtubule nucleation | 5 | 1 | TGME49_209970 | 20 | 28.61 | 36.55 | 0.0344<br>86 | 0.18598 | 1 |
| **GO:0019856** | pyrimidine nucleobase biosynthetic process | 5 | 1 | TGME49_215260 | 20 | 28.61 | 36.55 | 0.0344<br>86 | 0.18598 | 1 |
| **GO:0009190** | cyclic nucleotide biosynthetic process | 5 | 1 | TGME49_289170 | 20 | 28.61 | 36.55 | 0.0344<br>86 | 0.18598 | 1 |
| **GO:0009187** | cyclic nucleotide metabolic process | 5 | 1 | TGME49_289170 | 20 | 28.61 | 36.55 | 0.0344<br>86 | 0.18598 | 1 |
| **GO:0009163** | nucleoside biosynthetic process | 5 | 1 | TGME49_316100 | 20 | 28.61 | 36.55 | 0.0344<br>86 | 0.18598 | 1 |
| **GO:0006206** | pyrimidine nucleobase metabolic process | 5 | 1 | TGME49_215260 | 20 | 28.61 | 36.55 | 0.0344<br>86 | 0.18598 | 1 |
| **GO:0031109** | microtubule polymerization or depolymerization | 5 | 1 | TGME49_209970 | 20 | 28.61 | 36.55 | 0.0344<br>86 | 0.18598 | 1 |
| **GO:0009119** | ribonucleoside metabolic process | 6 | 1 | TGME49_316100 | 16.7 | 23.84 | 29.24 | 0.0412<br>44 | 0.207656 | 1 |
| **GO:0050896** | response to stimulus | 189 | 4 | TGME49_202540<br>TGME49_210960<br>TGME49_215060<br>TGME49_289170 | 2.1 | 3.03 | 3.35 | 0.0412<br>56 | 0.207656 | 1 |
| **GO:0034654** | nucleobase-containing compound biosynthetic process | 117 | 3 | TGME49_249560<br>TGME49_289170<br>TGME49_316100 | 2.6 | 3.67 | 4 | 0.0472<br>99 | 0.226289 | 1 |
| **GO:0097435** | supramolecular fiber organization | 7 | 1 | TGME49_209970 | 14.3 | 20.43 | 24.36 | 0.0479<br>55 | 0.226289 | 1 |

**Table S10.**
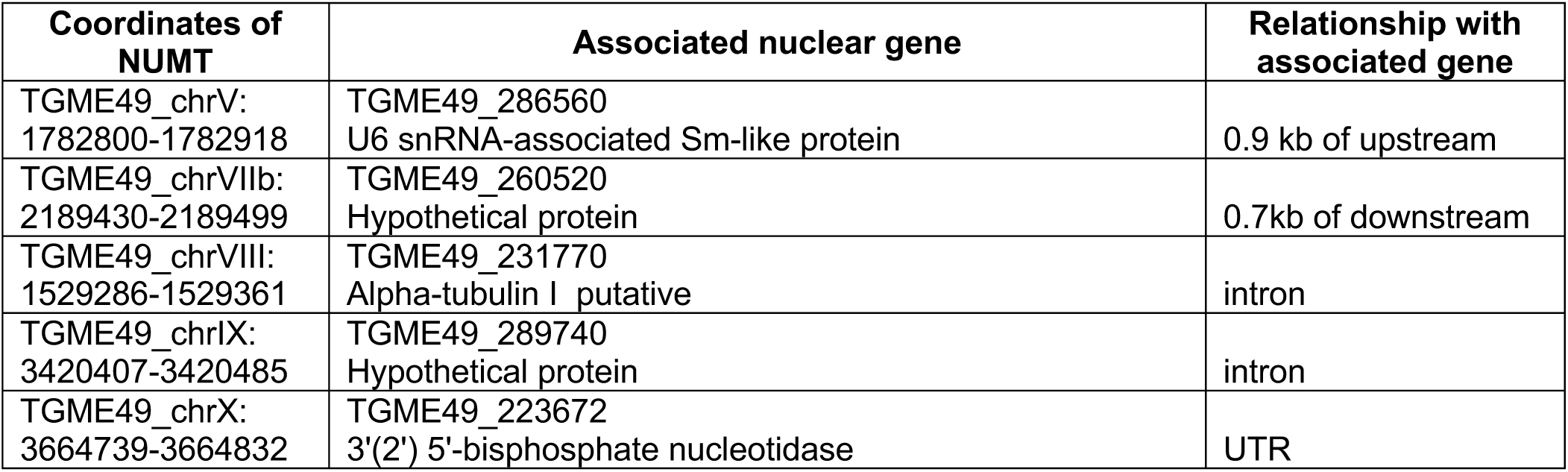
Orthologous NUMTs in *T. gondii* and *N. caninum*

**Table S11.**
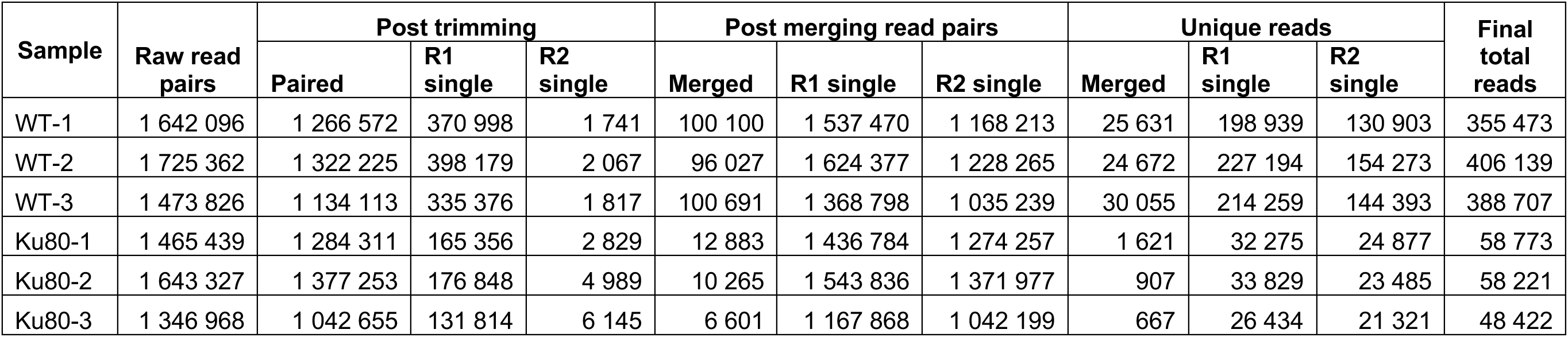
Amplicons sequencing read data summary

| Sample | Raw read pairs | Post trimming |  |  | Post merging read pairs |  |  | Unique reads |  |  | Final total reads |
| --- | --- | --- | --- | --- | --- | --- | --- | --- | --- | --- | --- |
|  |  | Paired | R1 single | R2 single | Merged | R1 single | R2 single | Merged | R1 single | R2 single |  |
| WT-1 | 1 642 096 | 1 266 572 | 370 998 | 1 741 | 100 100 | 1 537 470 | 1 168 213 | 25 631 | 198 939 | 130 903 | 355 473 |
| WT-2 | 1 725 362 | 1 322 225 | 398 179 | 2 067 | 96 027 | 1 624 377 | 1 228 265 | 24 672 | 227 194 | 154 273 | 406 139 |
| WT-3 | 1 473 826 | 1 134 113 | 335 376 | 1 817 | 100 691 | 1 368 798 | 1 035 239 | 30 055 | 214 259 | 144 393 | 388 707 |
| Ku80-1 | 1 465 439 | 1 284 311 | 165 356 | 2 829 | 12 883 | 1 436 784 | 1 274 257 | 1 621 | 32 275 | 24 877 | 58 773 |
| Ku80-2 | 1 643 327 | 1 377 253 | 176 848 | 4 989 | 10 265 | 1 543 836 | 1 371 977 | 907 | 33 829 | 23 485 | 58 221 |
| Ku80-3 | 1 346 968 | 1 042 655 | 131 814 | 6 145 | 6 601 | 1 167 868 | 1 042 199 | 667 | 26 434 | 21 321 | 48 422 |

**Table S12.**
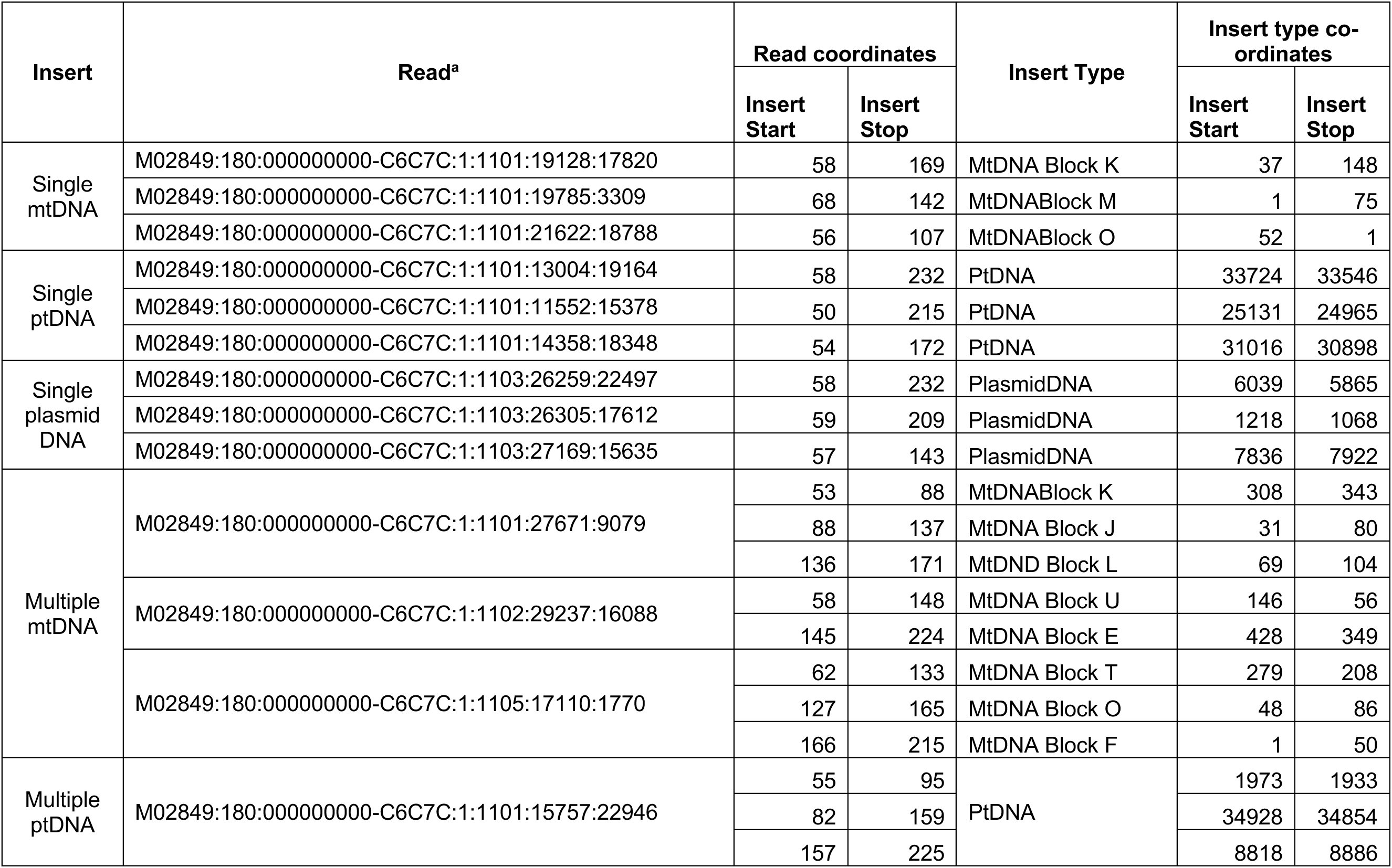
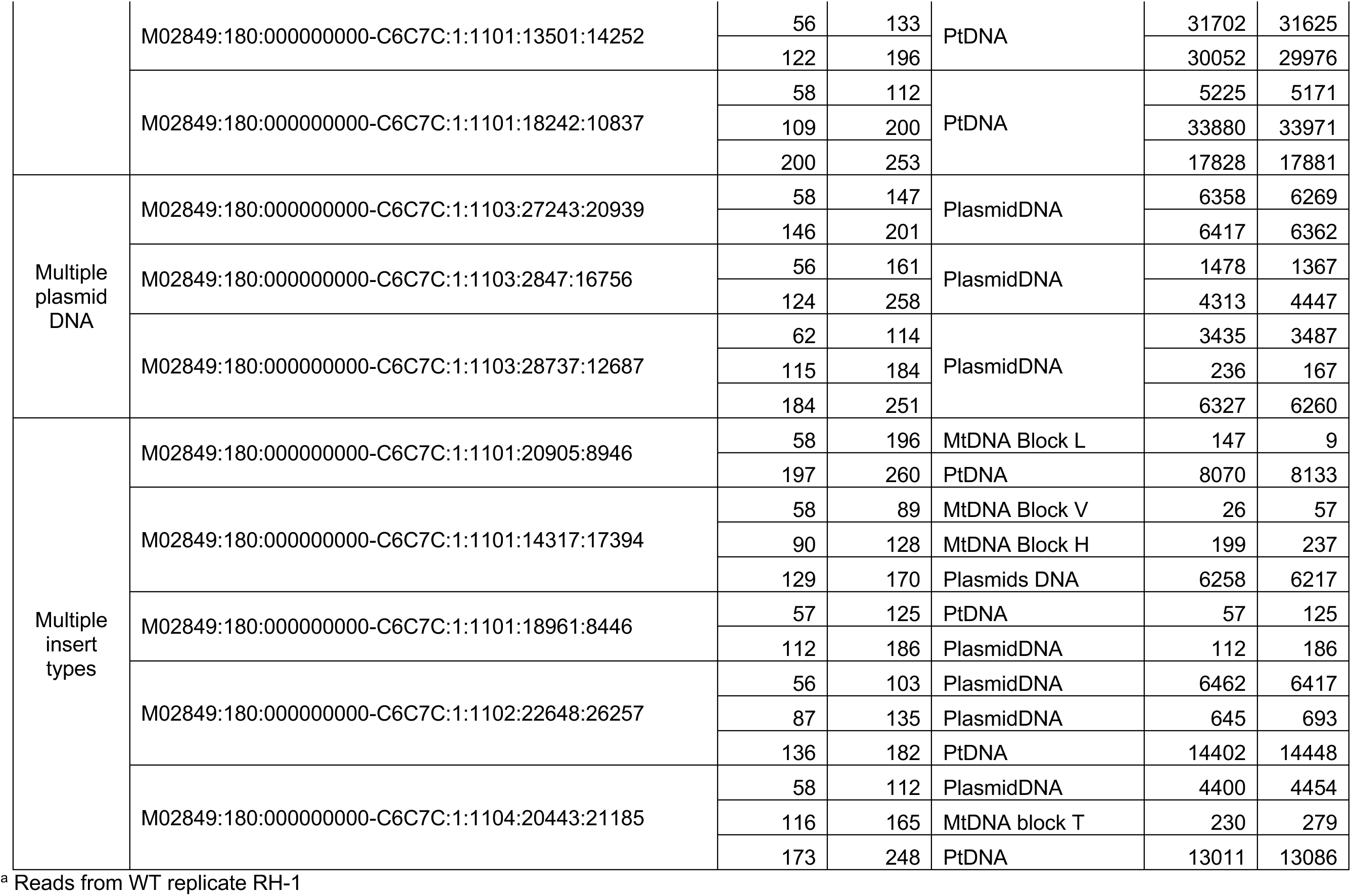
Examples of insertion types observed in amplicons from WT parasites

